# Structure of telomerase-bound CST with Polymerase α-Primase

**DOI:** 10.1101/2021.12.28.474374

**Authors:** Yao He, He Song, Henry Chan, Yaqiang Wang, Baocheng Liu, Lukas Susac, Z. Hong Zhou, Juli Feigon

## Abstract

Telomeres are the physical ends of linear chromosomes, composed of short repeating sequences (e.g. TTGGGG in *Tetrahymena* for the G-strand) of double-stranded DNA with a single-strand 3’-overhang of the G-strand and a group of proteins called shelterin^1,2^. Among these, TPP1 and POT1 associate with the 3’-overhang, with POT1 binding the G-strand^3^ and TPP1 recruiting telomerase via interaction with telomerase reverse transcriptase (TERT)^4^. The ends of the telomeric DNA are replicated and maintained by telomerase^5^, for the G-strand, and subsequently DNA Polymerase α-Primase^6,7^ (PolαPrim), for the C-strand^8^. PolαPrim is stimulated by CTC1–STN1–TEN1 (CST)^9–12^, but the structural basis of both PolαPrim and CST recruitment to telomere ends remains unknown. Here we report cryo-EM structures of *Tetrahymena* CST in the context of telomerase holoenzyme, both in the absence and presence of PolαPrim, as well as of PolαPrim alone. Ctc1 binds telomerase subunit p50, a TPP1 ortholog, on a flexible Ctc1 binding motif unveiled jointly by cryo-EM and NMR spectroscopy. PolαPrim subunits are arranged in a catalytically competent conformation, in contrast to previously reported autoinhibited conformation. Polymerase POLA1 binds Ctc1 and Stn1, and its interface with Ctc1 forms an entry port for G-strand DNA to the POLA1 active site. Together, we obtained a snapshot of four key players required for telomeric DNA synthesis in a single complex—telomerase core RNP, p50/TPP1, CST and PolαPrim—that provides unprecedented insights into CST and PolαPrim recruitment and handoff between G-strand and C-strand synthesis.

---

Synthesis of the G-strand at the ends of telomeres by telomerase is terminated by the heterotrimeric complex CST^12,13^. CST is essential for maintaining the telomere C-strand and also plays a role in overcoming genome-wide replication stress^9,14,15^. Like the shelterin proteins and telomerase, mutations in CST lead to telomere biology disorders^16^ including Coats Plus and dyskeratosis congenita^17^. CST small subunits STN1 and TEN1 are closely related to those in replication protein A (RPA)^18^, a singlestranded DNA binding protein involved in all aspects of DNA replication and repair, while the large subunit Ctc1/CTC1/Cdc13 is more diverse^19–24^. Structural and biochemical studies of CST proteins have suggested various stoichiometries, oligomerization states, and functions of subunits^19,20,24^. The only structure of complete CST^19^, from human, revealed a decameric architecture of heterotrimers in the presence of single-stranded telomeric (G-strand) DNA (sstDNA). CST is proposed to inhibit telomerase activity by physical interaction with shelterin proteins TPP1–POT1 at telomere ends^12,25,26^ and G-strand sequestration^12,13^, and it promotes C-strand fill-in by association with PolαPrim^27–30^, but structures of these interactions are lacking. PolαPrim is an unusual polymerase containing both primase and DNA polymerase subunits; the primase synthesizes an RNA primer on a DNA template and then hands off the duplex to the polymerase that initiates synthesis of a short DNA duplex^6,7^.

*Tetrahymena* telomerase holoenzyme comprises, in addition to its RNP catalytic core of TERT, telomerase RNA (TER) that provides the template for G-strand telomere repeat synthesis, and LARP7 assembly protein p65, several proteins that are orthologous to human proteins that only transiently associate with telomerase at telomeres^31,32^. These include p50, the structural and functional equivalent of TPP1 that recruits and activates telomerase^23,33,34^; Teb1, a subunit of a trimeric RPA related complex TEB^23^, that binds the sstDNA^35^ and together with p50 increases activity and processivity like its ortholog POT1^36–38^; and another trimeric RPA related complex p75–p45–p19 that has been identified as *Tetrahymena* Ctc1–Stn1–Ten1 (*Tt*CST)^23,24^. The constitutive association of these proteins with telomerase catalytic core makes *Tetrahymena* telomerase an ideal model system for elucidating details of the protein structures and interactions that regulate G-strand and C-strand synthesis^23,39,40^. Structural studies of *Tetrahymena* telomerase and PolαPrim described here show how monomeric *Tt*CST binds p50/TPP1 and PolαPrim on different interfaces to coordinate G-strand termination and C-strand fill-in synthesis, and suggest commonalities with interactions at human telomeres.

## Overall structure of telomerase bound CST

While cryo-EM studies of *Tetrahymena* telomerase have provided high-resolution structures of the RNP catalytic core, TEB heterotrimer, and p50 OB^40^, the dynamic positioning of *Tt*CST has limited its structure modeling to date. Here, we combined three previously reported datasets of *Tetrahymena* telomerase bound to sstDNA^40^ and conducted focused classification on *Tt*CST followed by refinement of the holoenzyme to obtain a reconstruction with an overall resolution of 3.5 Å (Fig. 1a, Extended Data Fig. 1, Extended Data Table 1). For the model of *Tt*CST, Stn1–Ten1 crystal structures^23,24^ were rigid body fit into the density and manually refined with little change, and Ctc1 was built *de novo* (Fig. 1b, Extended Data Fig. 2). Modeling of the N-terminal domain of Ctc1, which has lower resolution (Extended Data Fig. 1c), was facilitated by using information derived from NMR data on secondary structure elements and inter-β-strand NOEs (see Methods) (Extended Data Fig. 3). Ctc1, whose domain structure was not previously established, comprises three OBs (OB-A, -B, and -C) connected by structured linkers that stabilize the rigid pairwise interactions between the domains (Fig. 1c, d). Ctc1 OB-C has a C-shaped cleft and Zn-ribbon motif (Fig. 1c) typical of the C-terminal OB of the large subunit of RPA^18^ and related complexes, including mammalian CTC1^19^, POT1^37,38^, and *Tt*Teb1^35,40^. Ctc1 OB-C forms a heterotrimer with Stn1 and Ten1 OB that is stabilized by an intermolecular three-helix bundle (Fig. 1c, e) and by Ten1–Stn1 and Stn1–Ctc1 OB interactions (Extended Data Fig. 2c-g). The tandem <u>w</u>inged <u>h</u>elix-turn-helix (WH-WH) domain of Stn1 is connected to OB by a flexible linker and is not visible in the cryo-EM map, consistent with its multi-positioning shown by negative-stain EM^23^. Overall, our structure of monomeric *Tt*CST strongly suggests its origin from RPA and establishes the domain structure of the least conserved subunit Ctc1.

**Fig. 1:**
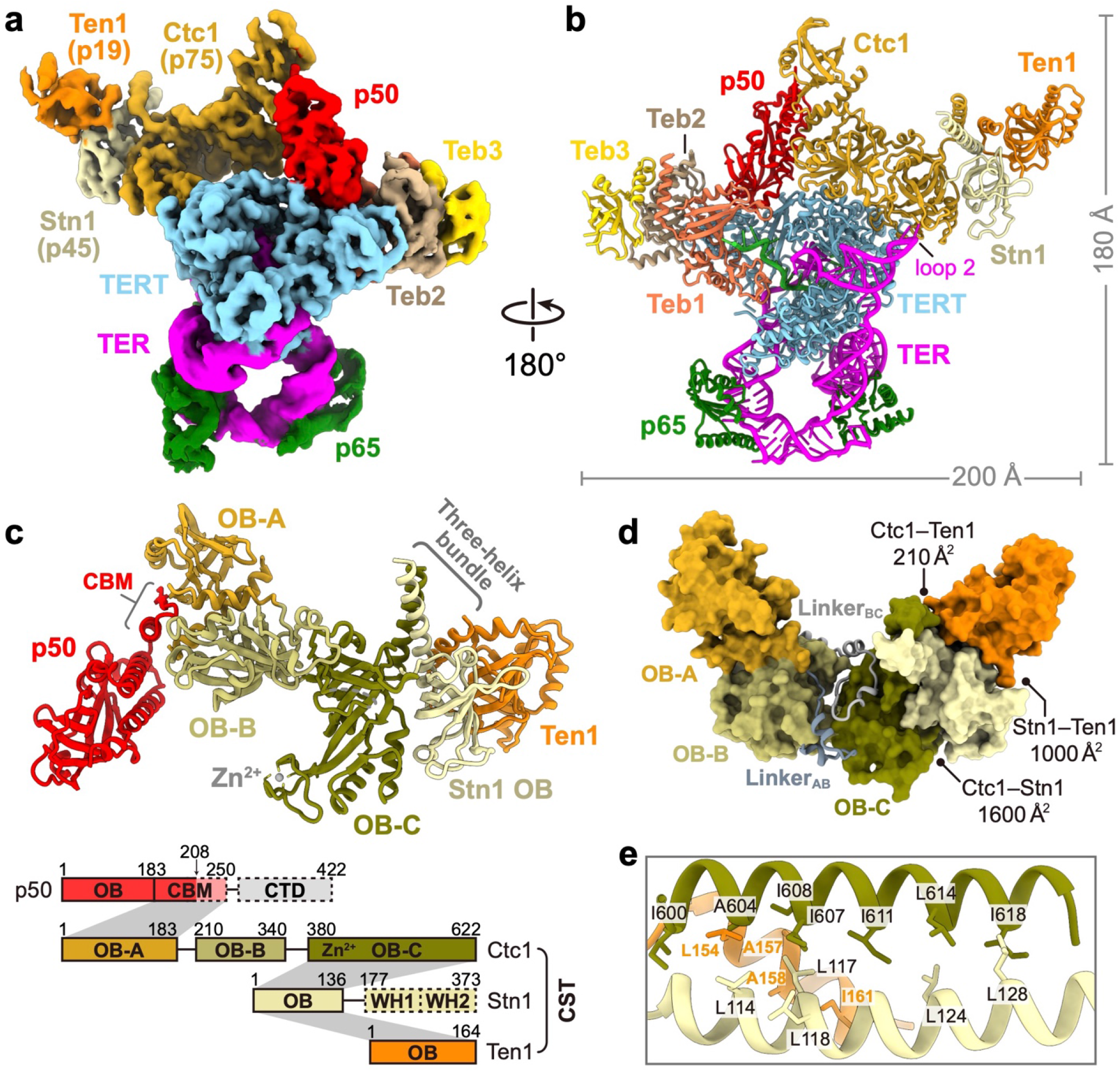
Cryo-EM structure of *Tt*CST in telomerase holoenzyme. **a,** Cryo-EM map of telomerase holoenzyme at 3.5 Å resolution. **b,** Ribbon representation of the model of telomerase holoenzyme, viewed after a 180° rotation from **a**. The proteins, TER and DNA are color-coded as indicated. **c,** Structure and schematic of *Tt*CST with p50. The three OB domains (OB-A, -B, and -C) of Ctc1 are colored individually as indicated. In the schematic, invisible regions in the cryo-EM map are shown as dashed boxes. Intermolecular interactions between proteins are indicated as gray shading. OB, oligonucleotide/oligosaccharide-binding fold domain; WH, winged-helix domain; CBM, CST binding motif; CTD, C-terminal domain; Zn^2+^, Zn-ribbon motif. **d,** Surface representation of *Tt*CST structure. Buried surface areas in the interfaces between *Tt*CST subunits are indicated. The two structured linkers between Ctc1 OB domains are shown as ribbon. **e,** Zoom-in view of the hydrophobic interface of *Tt*CST intermolecular three-helix bundle.

## Flexible interface between Ctc1 and p50

p50 has an N-terminal OB and a C-terminal domain that is invisible in the cryo-EM map (Fig. 1a). p50/TPP1 OB interacts with telomerase on TERT TEN and TRAP domains^39–42^, but how TPP1–POT1 interacts with CST is unknown^25,26^. We find that *Tt*CST is anchored on p50 via Ctc1 OB-A (Fig. 1c). In the structure, *Tt*CST is positioned across the top of the TERT ring (Fig. 1b), and stabilized in this predominant conformation by additional interactions between Ctc1 and TERT–TER catalytic core (Extended Data Fig. 2h-k). However, these are not stable interactions, as other conformations resolved by 3D classification show *Tt*CST hinged away from TERT (Fig. 2a, Extended Data Fig. 1a). In the cryo-EM map, a previously uncharacterized density of p50 protrudes from its OB C-terminus into Ctc1 OB-A (Extended Data Fig. 1g). p50 residues 185-208 were built against the density as helix α5 and strand β7, the latter of which forms an extended β sheet with β1-β4-β5’ of Ctc1 OB-A (Fig. 2b). However, previous biochemical studies showed that p50 C-terminal truncation at residue 213 almost abrogates binding with Ctc1, while truncation at residue 252 showed binding with Ctc1 comparable to full-length protein^43^. We therefore investigated whether p50 residues between 208-255 might be contributing to the binding interface with Ctc1. We made a series of p50 peptides and monitored their interaction with Ctc1 OB-A by NMR (Extended Data Fig. 4). ^1^H-^15^N HSQC spectra show that optimal binding requires residues 228-250 (Fig. 2c, Extended Data Fig. 4b). This peptide forms a 1:1 complex with Ctc1 OB-A that is in slow exchange on the NMR timescale indicating slow off-rate (Extended Data Fig. 4c). Talos+ (ref^44^) secondary structure scores, CS-Rosetta^45^ modeling, and chemical shift mapping indicate that p50 peptide residues 228-241 form a β-hairpin that interacts with the Ctc1 OB-A β-barrel near β1-β2 linker, β4, and β5 (Fig. 2d, Extended Data Fig. 4e-g). Together, these cryo-EM and NMR data define a Ctc1 binding motif (CBM) adjacent to p50 OB that tightly associates with Ctc1 but allows hinging movement of the entire CST complex on p50.

**Fig. 2:**
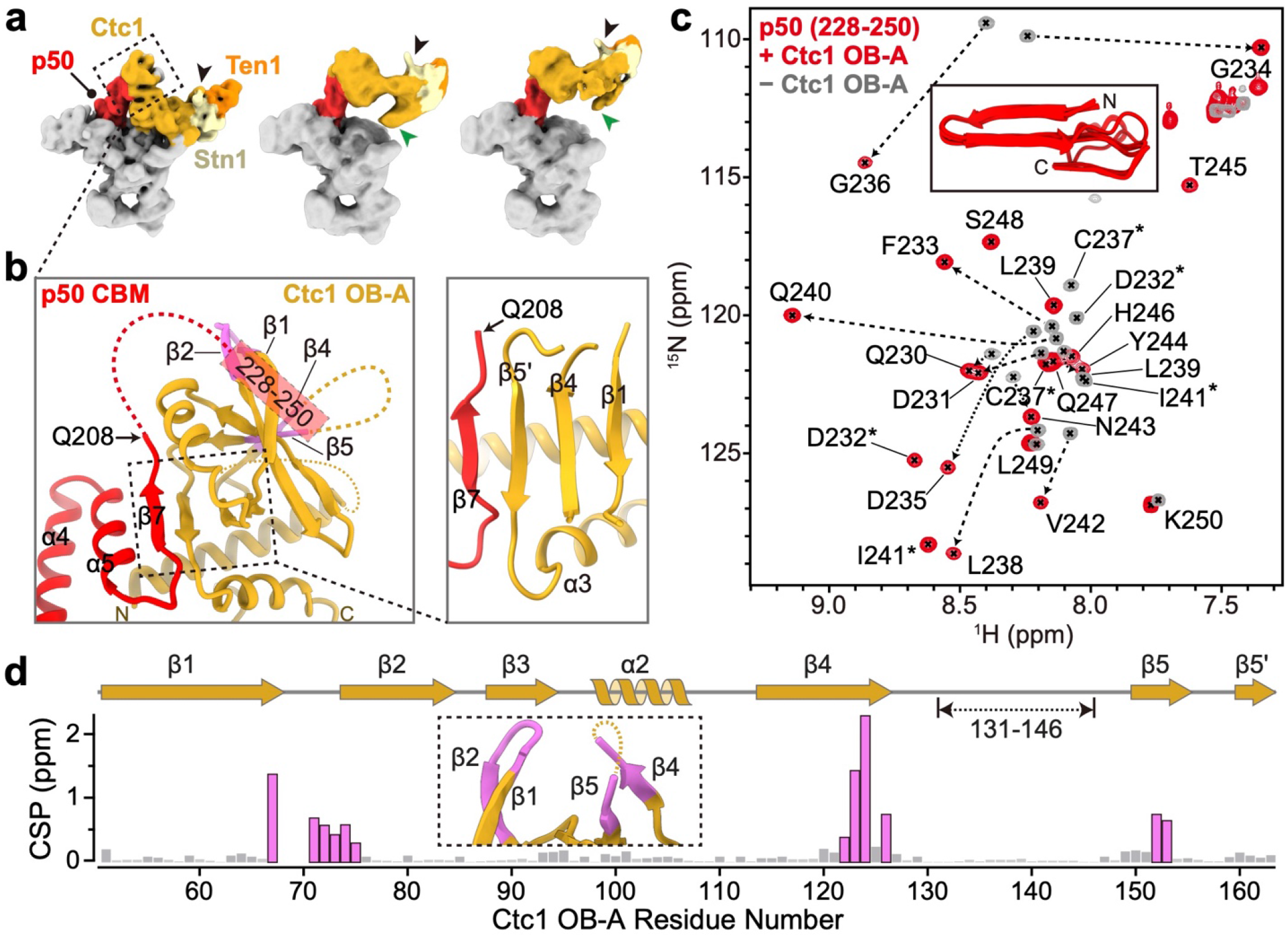
Interface between *Tt*CST and p50. **a,** Cryo-EM maps of telomerase holoenzyme with *Tt*CST at different positions. *Tt*CST subunits and p50 are colored as indicated. The three-helix bundle and Zn-ribbon motif are labeled with black and green arrows, respectively. Cryo-EM maps are low-pass filtered to similar resolution for comparison. **b,** Interactions between Ctc1 OB-A and p50 CBM. Unmodeled regions in the cryo-EM structure are shown as dashed lines. **c,** ^1^H-^15^N HSQC spectra of ^15^N-labelled p50 peptide (residues 228-250) with (red) and without (gray) Ctc1 OB-A. NMR signals from the same residues are connected with dashed arrows or labeled with asterisks. Inset shows CS-Rosetta models of p50 peptide in the presence of Ctc1 OB-A. **d,** Chemical shift perturbation (CSP) index of ^15^N-labelled Ctc1 OB-A upon binding p50 peptide. Residues with CSP over 0.25 ppm are highlighted in magenta, and their locations on the cryo-EM structure are shown in an inserted panel and on **b**.

## Overall structure of PolαPrim with CST

A defining feature of CST is its ability to recruit PolαPrim for C-strand synthesis; however, the interface between PolαPrim and CST has not been structurally characterized in any organism^9,27–30,46^. To both verify that p75–p45–p19 is functionally *Tt*CST and define the mechanism of PolαPrim recruitment, we assembled *Tetrahymena* telomerase–PolαPrim complex using endogenously expressed telomerase and recombinant PolαPrim in the presence of sstDNA d(GTTGGG)_10_, and determined its cryo-EM structure (Fig. 3a-c, Extended Data Fig. 5, 6). In the telomerase holoenzyme PolαPrim binds *Tt*CST in the absence or presence of sstDNA (Fig. 3c). Since the entire *Tt*CST–PolαPrim complex was flexibly positioned relative to p50, as seen for *Tt*CST alone, the *Tt*CST–PolαPrim and telomerase RNP core– TEB–p50 complexes were processed separately to obtain 3D reconstructions of 4.2 Å and 2.9 Å resolution, respectively (Fig. 3a, Extended Data Fig. 6, Extended Data Table 1). The modeled structures are the first of CST-bound PolαPrim and the highest resolution structure of telomerase RNP core to date (Fig. 3b, Extended Data Fig. 6f). Modeling of *Tt*CST in the complex by initial rigid body fitting of the structure determined in the absence of PolαPrim revealed the presence of additional density on Ctc1 OB-B and OB-C that could be fit with the crystal structure of Stn1 WH-WH^23,24^ (Fig. 3b, Extended Data Fig. 6f). Binding of PolαPrim to *Tt*CST displaces Ctc1 from its stable position across the top of the TERT ring (compare Fig. 3a, b to Fig. 1a, b), and instead POLA1 is positioned near TER loop 2. The Stn1 WH-WH binding site on Ctc1 would also be occluded in the stable conformation of *Tt*CST on p50–TERT in the absence of PolαPrim.

**Fig. 3:**
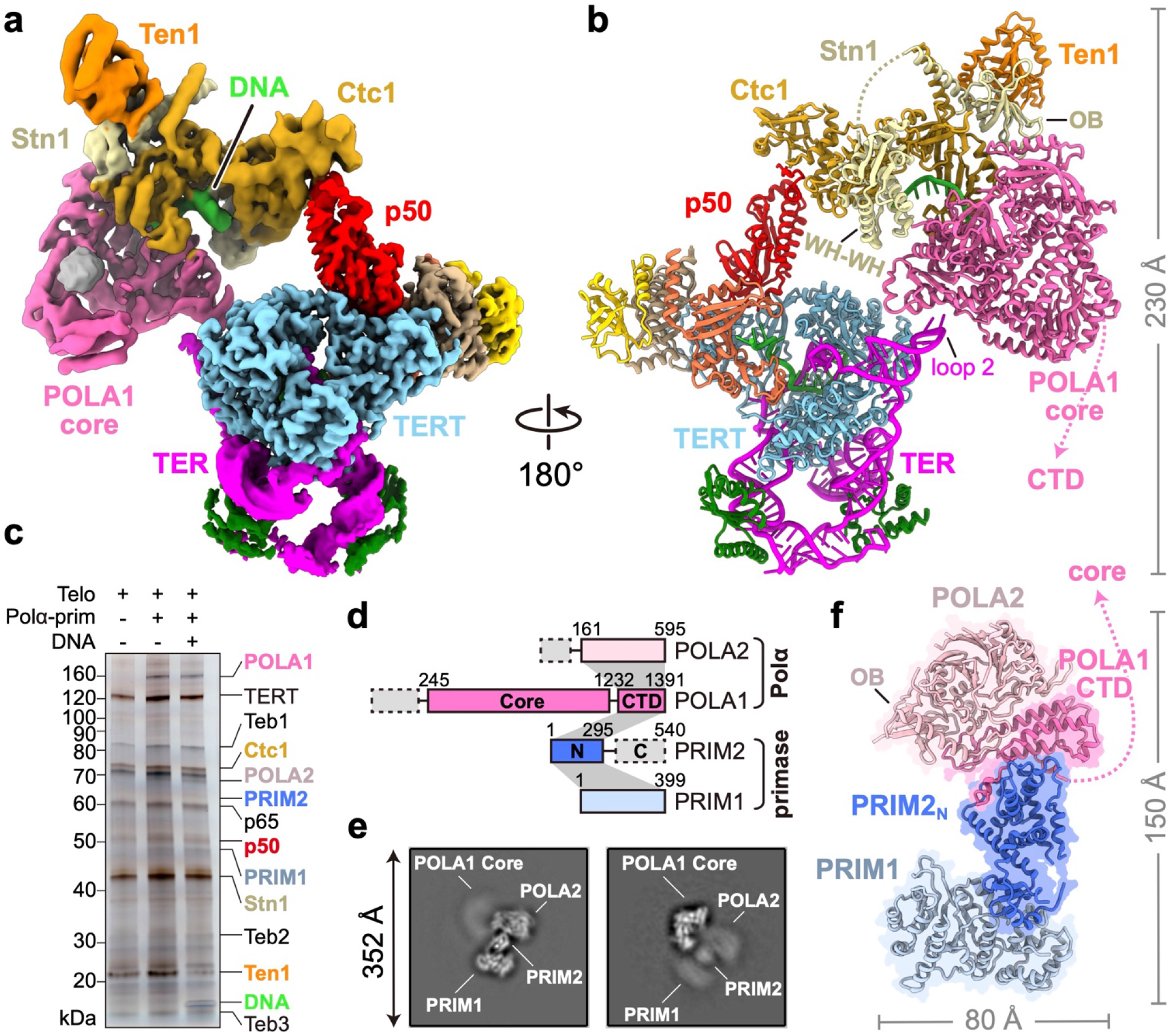
Structure of *Tetrahymena* telomerase holoenzyme in complex PolαPrim. **a,** Composite map of the complex generated with focused refined cryo-EM maps (Extended Data Fig. 6a). **b,** Atomic model of the complex. The unstructured linker between Stn1 OB and WH-WH domains is shown as dashed line. **c,** Silver-stained SDS-PAGE gel of affinity purified samples shown assembly of the complex with or without sstDNA. **d,** Structure-based diagram of PolαPrim. Intermolecular interactions between subunits are indicated as gray shading. **e,** Representative 2D-class averages of PolαPrim. **f,** Cryo-EM structure of PolαPrim. The flexible linker between POLA1_core_ and POLA1_CTD_ is shown as dashed line.

PolαPrim comprises two polymerase (POLA1 and POLA2) and two primase (PRIM1 and PRIM2) subunits^6,7^ (Fig. 3d). The presence of all four subunits in the complex was confirmed by silver-stain SDS-PAGE (Fig. 3c) and negative-stain EM 2D classification analysis (Extended Data Fig. 5f). However, only the catalytic POLA1_core_ was well resolved in the cryo-EM map (Fig. 3a, d). The rest of PolαPrim appears as a fuzzy density connected to POLA1_core_ opposite the interface with Ctc1 (Extended Data Fig. 5f). Therefore, we also investigated the cryo-EM structure of PolαPrim alone, and obtained a 4.0-4.3 Å resolution structure for POLA2–POLA1_CTD_–PRIM2_N_–PRIM1 (Fig. 3e, f, Extended Data Fig. 7, Extended Data Table 1). 2D class averages of PolαPrim show that POLA2–POLA1_CTD_– PRIM2_N_–PRIM1 forms a platform that holds POLA1_core_ at various positions (Fig. 3e). Initial models of PolαPrim subunits generated using AlphaFold2^47^ were rigid-body fit into corresponding densities and manually adjusted. The structures of the individual subunits are highly similar to those of human PolαPrim^48–51^ (Extended Data Fig. 8a, b). However, POLA2-POLA1_CTD_ and PRIM1 that are located on either end of the platform can apparently rotate relative to each other, with PRIM2_N_ as the pivot (Extended Data Fig. 8b). PRIM2_C_, which specifically interacts with and coordinates RNA–DNA duplex translocation from the active site on PRIM1 to the active site on POLA1_core_ (ref^52^), was not observed during cryo-EM data processing, suggesting its dynamic positioning. Such flexible organization of PolαPrim would allow for the large-scale domain movements expected for the switch from RNA primer to C-strand DNA synthesis^7^. Previous structures of human PolαPrim determined by X-ray crystallography and cryo-EM of a crosslinked sample are in an autoinhibited conformation (Extended Data Fig. 8c) with the active site on POLA1_core_ sterically blocked by POLA1_CTD_ and POLA2 for DNA entry^48,49^. Here, our studies provide the first structures of a PolαPrim compatible with activity, and they establish its direct interactions with *Tt*CST, as described in detail next.

## CST interaction with POLA1 and sstDNA

POLA1_core_ comprises an N-terminal domain (NTD) that brackets a catalytically dead exonuclease (Exo), and a C-terminal DNA polymerase that contains palm, fingers, and thumb domains^7^ (Fig. 4a, b). All elements of POLA1_core_ except the tip of the thumb are well defined in the cryo-EM map. POLA1_core_ interacts with CST via its Exo and C-terminal region of NTD, which positions its active site ~40 Å from the interface (Fig. 4a). CST interacts with POLA1_core_ via Ctc1 OB-C and Stn1 (Fig. 4b-d). On Ctc1 OB-C, the conserved Zn-ribbon motif interacts primarily with conserved Exo β11-β12 hairpin (Fig. 4c, Extended Data Fig. 8d). Ctc1 helix α14 and Stn1 OB β1-β2 and β3-β4 loops form a binding pocket that accommodates POLA1_core_ NTD helix α19 (residues 731-748) with charge complementarity to its side chains (Fig. 4c, Extended Data Fig. 8e-g). Helix α19 is a flexible loop in all other structures of PolαPrim^48–51,53^, and apparently becomes structured only on binding Ctc1–Stn1 (Extended Data Fig. 8e), indicating the importance of this interaction for CST binding. Structure-based sequence alignment suggests conservation of these interfaces on POLA1_core_ across a wide-range of species (Extended Data Fig. 8g). Behind this interface, the three helices of Stn1 WH2 are inserted into a gap between Ctc1 OB-B and POLA1 Exo (Fig. 4d).

**Fig. 4:**
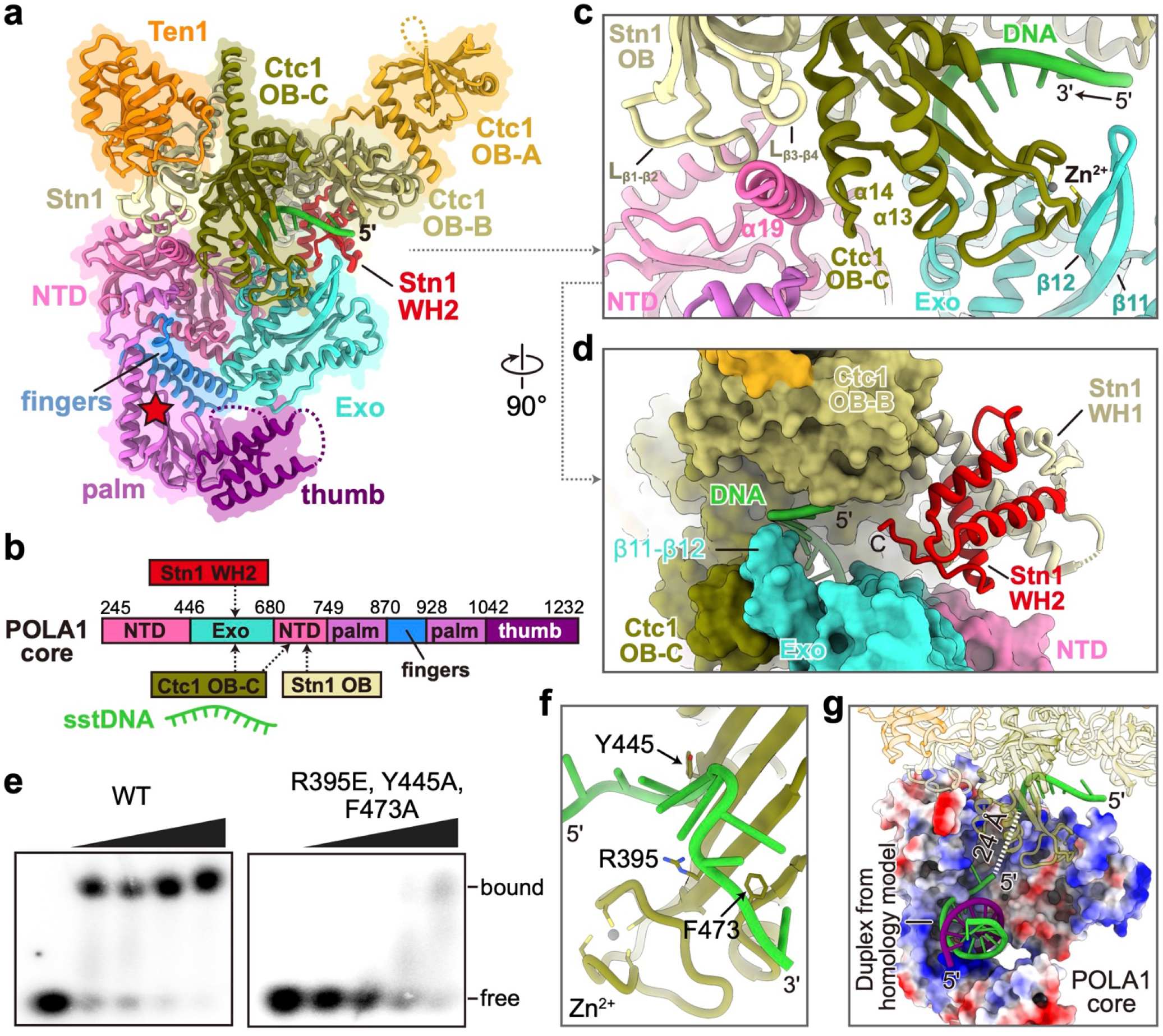
Interface between *Tt*CST and POLA1_core_. **a,** Ribbon representation of *Tt*CST and POLA1_core_ with individual protein/domain/motif colored as indicated. Location of POLA1 active site is shown as a red star. NTD, N-terminal domain; Exo, exonuclease domain. **b,** Structure-based diagram of POLA1_core_ and interactions with *Tt*CST. **c, d,** Zoom-in views of the interface between POLA1_core_ and *Tt*CST with sstDNA shown from perpendicular directions. **e,** EMSA of d(GTTGGG)_5_ DNA binding by wild-type (WT) and Ctc1 mutant *Tt*CST. Wedges indicate two-fold dilution of *Tt*CST starting at 2.5 μM. The first lane of each gel is a *Tt*CST-free control. **f,** sstDNA binding site on Ctc1 OB-C. Residues substituted for EMSA are shown as sticks. **g,** Zoom-in view of POLA1_core_ (electrostatic surface) with a DNA duplex modeled based on human POLA1 structure (PDB 5IUD) and path of the sstDNA in the channel shown as dashed line. The template and product strands in the duplex are colored in green (G-strand) and purple (C-strand), respectively.

Cryo-EM density for ~10 nucleotides of sstDNA is observed on Ctc1 OB-C across the C-shaped binding cleft near the Zn-ribbon (Fig. 4a, c). Substitution of three conserved residues R395E, Y445A and F473A on the binding surface almost abolishes sstDNA binding to purified *Tt*CST, as determined by electromobility shift assays (EMSA) with d(GTTGGG)_5_ (Fig. 4e, f). The sstDNA extends 5’!3’ into an entry port, formed by POLA1_core_ NTD and Exo and Ctc1 OB-C (Extended Data Fig. 8h), of a highly basic channel that leads to the active site of POLA1_core_, where the primer–sstDNA duplex would bind (Fig. 4g). Since sstDNA is added in excess during the purification, both telomerase and *Tt*CST can bind separate sstDNA strands. No density for sstDNA is observed on *Tt*CST in the absence of PolαPrim, perhaps due to dynamics and/or steric occlusion by interaction of Ctc1 with TERT in the predominant conformation (Fig. 1b, Extended Data Fig. 2h-k).

The structure shows that POLA1_core_ Exo and NTD form an extensive interface with *Tt*CST involving Ctc1 OB-C, Stn1 OB, and Stn1 WH1, which is otherwise flexibly tethered to Stn1 OB in the absence of PolαPrim. The sstDNA on Ctc1 OB-C appears positioned for entry into a template (G-strand) binding tunnel on POLA1_core_, and there is weak density within the tunnel that we attribute to sstDNA (Extended Data Fig. 8i, j), suggesting the possibility that this structure has captured the polymerase mode after handoff from primase. Although no DNA–RNA or DNA–DNA duplex is present, there is unassigned density between POLA1_core_ palm and thumb (Fig. 3a) that fits the dimensions of a G-quadruplex formed by four *Tetrahymena* telomere repeats (Extended Data Fig. 8j). We propose that the G-quadruplex may have serendipitously trapped the PolαPrim in an incipient DNA polymerization state.

## Comparison to human CST

While *Tetrahymena* Stn1 and Ten1 have the same domain structure as human STN1 and TEN1, respectively, human CTC1 (hCTC1) is much larger than *Tetrahymena* Ctc1, with seven OBs (OB-A through OB-G)^19^ (Fig. 5a, b, Extended Data Fig. 9a) that may have arisen from a gene duplication of RPA70 (e.g. OB-N,A,B,N,A,B,C). A DALI^54^ search of Ctc1 against all proteins in the PDB found the highest structural similarity with hCTC1 (Z-score 14.5). For the individual domains, Ctc1 OB-B and OB-C are most similar to hCTC1 OB-F and OB-G, respectively (Extended Data Fig. 9b, c). Cryo-EM study of hCST with bound sstDNA revealed a decameric structure with D5 symmetry^19^. Comparing the hCST monomer extracted from the decamer to *Tt*CST shows Ten1/TEN1, Stn1 OB/STN1 OB, and Ctc1 OB-A,B,C/CTC1 OB-E,F,G are positioned similarly (Fig. 5a, b). However, Stn1 WH-WH, which is only visible in the PolαPrim bound *Tt*CST structure, is positioned on Ctc1 OB-B and OB-C (Fig. 5a, Extended Data Fig. 9d), while in hCST STN1 WH-WH is positioned on OB-E and sticks out from the decamer in what has been called the ‘Arm’ conformation^19^ (Fig. 5b, c). Intriguingly, low resolution cryo-EM densities of monomeric hCST revealed an additional conformation^19^, called ‘Head’, where hSTN1 WH-WH occupies a position apparently close to that observed in *Tt*CST bound PolαPrim (Fig. 5c). In our structure, Stn1 WH-WH in this position forms part of the interface with PolαPrim (Fig. 4d). If hCST binds PolαPrim in a similar manner to *Tt*CST, it could only bind as monomer since the binding interface would be occluded by intermonomer interactions in the hCST decamer.

**Fig. 5:**
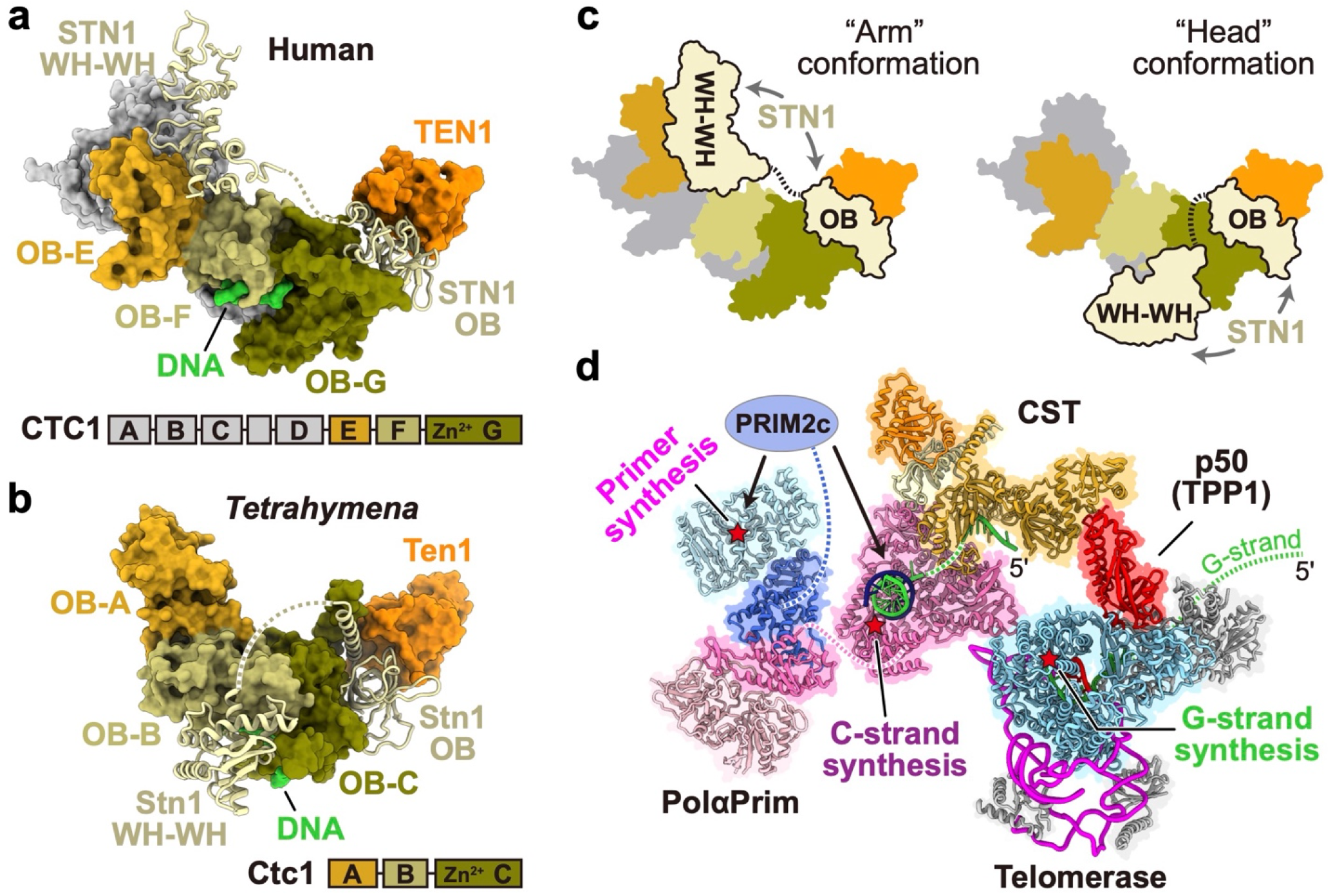
Structural comparison of *Tt*CST and hCST. **a, b,** Surface representation of *Tt*CST (a) and hCST (PDB 6W6W) (b) with Stn1/STN1 shown as ribbons. Corresponding subunits/domains are colored the same as indicated. **c,** Cartoon illustrations of hCST with STN1 WH-WH in “arm” and “head” conformations^19^. The linker between STN1 OB and WH-WH is shown as dashed line. **d,** Model of *Tetrahymena* telomerase holoenzyme in complex with PolαPrim. The DNA duplex on POLA1_core_ is modeled based on a homology model of human POLA1_core_ (PDB 5IUD). Position of the POLA2– POLA1_CTD_–PRIM2_N_–PRIM1 platform relative to POLA1_core_ is based on a low-resolution cryo-EM map in Extended Data Fig. 7a. Telomeric DNA G-strand and C-strand are colored green and purple, respectively. PRIM2_C_ is shown as an oval connecting to PRIM2_N_. Active sites on TERT, POLA1 and PRIM1 for the synthesis of G-strand, C-strand and RNA primer, respectively, are denoted by red stars.

Different but adjacent binding sites for sstDNA are observed for equivalent regions on hCST decamer^19^ vs *Tt*CST (Fig. 5a, b, Extended Data Fig. 9e). For hCST, the four nucleotides visible in the decameric structure interact with hCTC1 OB-F (Extended Data Fig. 9e), with two also interacting with OB-G^19^. Comparison of Ctc1 OB-B to CTC1 OB-F shows sequence similarity for the interacting residues on the sstDNA binding cleft (Extended Data Fig. 9f). Substitution of two sets of these conserved residues moderately decreases sstDNA binding as assayed by EMSA (Extended Data Fig. 9g-i). Together, these comparisons suggest hCTC1 C-terminal OB-E, -F, -G may interact with POLA1 and sstDNA in a similar manner to Ctc1 OB-A, -B, -C, and that PolαPrim interaction could only be accommodated on monomeric CST.

## Coordinated synthesis of G- and C-strands

Co-ordinated synthesis of telomeric G- and C-strands is orchestrated by interactions between telomerase, shelterin proteins TPP1 and POT1, sstDNA, CST, and PolαPrim, whose molecular details have been largely undefined. Taking advantage of the constitutive association of p50/TPP1, Teb1/POT1, and *Tt*CST with *Tetrahymena* telomerase along with previous structural studies of telomerase we determined structures and interfaces between all of these components. *Tt*CST Ctc1 C-terminal domain OB-C and Stn1 bind to PolαPrim POLA1 and Ctc1 N-terminal domain OB-A binds p50/TPP1 (Fig. 5d). p50/TPP1 OB in turn binds to TERT TEN–TRAP domains, constitutively in *Tetrahymena* but transiently to recruit telomerase to telomeres in humans^2,42^. Biochemical studies have shown that CST binds G-strands released from telomerase during telomere repeat synthesis^12,13^. Our structures show that *Tt*CST binds p50 on a flexible hinge (CBM), placing it in proximity to where the 3’-end of sstDNA released from the TER template would be. Teb1, that also interacts with p50, could contribute by initially maintaining a hold on the 5’ exiting DNA, until it binds *Tt*CST. C-strand synthesis requires that the released 3’-end of the G-strand enters the active site of the primase, where a short RNA primer is synthesized. PRIM2c binds the RNA–DNA duplex and hands it to the active site of POLA1 (Fig. 5d). Our structure explains how CST enhances the activity of POLA1, by binding the G-strand and feeding it into the entry port for the POLA1 template channel (Fig. 4g), thereby unwinding and preventing the reformation of any G-quadruplexes^55^. The autoinhibited conformation of PolαPrim^48,49^ would be occluded in the CST complex, possibly explaining how CST could enhance the primase-to-polymerase switch. It is less clear how CST binding could activate PRIM1 for primer synthesis, consistent with proposals for a large conformational switch between priming and polymerization steps. Here we have captured a pre-DNA polymerization step in a recruitment complex with *Tetrahymena* telomerase, linking G-strand and C-strand synthesis.

## Methods

### *Tetrahymena* PolαPrim cloning and expression

*Tetrahymena* PolαPrim complexes were expressed using the Bac-to-Bac system (Thermo Fisher Scientific) in *Sf9* cells. Briefly, cDNAs encoding *Tetrahymena* POLA1 (Accession number: Q23AJ0), POLA2 (I7MAE1), PRIM1 (Q24HY6) and PRIM2 (Q246C7) were chemically synthesized and purchased from IDT (Integrated DNA Technologies, Inc.). To co-express the POLA1–POLA2 complex, POLA1 and POLA2 cDNAs were cloned into a pFastBacDual vector (Thermo Fisher Scientific), under the polyhedrin promoter and the p10 promoter, respectively. The POLA1 has an N-terminal hexa-histidine-TEV (His_6_-TEV) tag, in which TEV is a tobacco etch virus protease cleavage site. To co-express the POLA1–POLA2–PRIM1–PRIM2 complex, PRIM1 and PRIM2 cDNAs were cloned into a separate pFastBacDual vector. The expression vectors were used to make baculoviruses based on the established protocol for Bac-to-Bac system (Thermo Fisher Scientific). *Sf9* cells (2.0 × 10^6^/ml) were transfected with viruses using a multiplicity of infection (MOI) of 3 at 27°C in SF-900™ II SFM media (Thermo Fisher Scientific). The cells were harvested 48 h after infection and stored at −80°C until purification.

### *Tetrahymena* CST–p50 cloning and expression

*Tetrahymena* CST–p50 complex was expressed in insect cells. Briefly, cDNAs encoding p50 (D2CVN8), Ctc1/p75 (A0PGB2), Stn1/p45 (Q6JXI5) and Ten1/p19 (D2CVN7) were chemically synthesized and purchased from IDT (Integrated DNA Technologies, Inc.). The Ctc1 and p50 cDNAs were cloned into a pFastBacDual vector (Thermo Fisher Scientific), with a His_6_-TEV tag fused onto the N-terminal of Ctc1. The Stn1 and Ten1 cDNAs were cloned into a separate pFastBacDual vector for baculoviruses expression. *Sf9* cells (2.1 × 10^6^/ml) were transfected with viruses using a multiplicity of infection (MOI) of 3 at 27°C in SF-900™ II SFM media (Thermo Fisher Scientific). The cells were harvested 48 h after infection and stored at −80°C until purification.

### Purification of *Tetrahymena* PolαPrim and CST–p50 complexes

The purification steps for both POLA1–POLA2–PRIM1–PRIM2 and p50–Ctc1–Stn1–Ten1 were performed at 4°C using an AKTA chromatography system with prepacked columns (GE Healthcare), following the same protocol. Cells were suspended in buffer A [30 mM Tris-HCl (pH 7.5), 200 mM NaCl, 10% (v/v) glycerol, 1mM dithiothreitol (DTT), and 25 mM imidazole] supplemented with protease inhibitor cocktail (Sigma), lysed by sonication, and centrifuged at 34,000×*g* for one hour. The supernatant was applied onto a 5-ml HisTrap HP column pre-equilibrated in buffer A. The column was washed with buffer A and the complex was eluted with buffer B [30 mM Tris-HCl (pH 7.5), 1 M NaCl, 10% (v/v) glycerol, 1mM DTT, and 400 mM imidazole]. The protein complex was digested overnight with 0.2 mg/ml TEV protease and buffer-exchanged to buffer A. The digest was applied onto a 5-ml HisTrap HP column pre-equilibrated in buffer A. The target complex was isolated in the column flow-through, concentrated to 10 ml, and then applied onto a Superdex 200 gel filtration column pre-equilibrated in buffer C [25 mM Tris-HCl (pH 7.5), 150 mM NaCl, and 1mM DTT]. The complex was collected from peak fractions and analyzed by SDS polyacrylamide gel electrophoresis.

### NMR sample preparation

cDNAs of Ctc1 OB-A (residues 1-183) and p50 peptides (Extended Data Fig. 4a) were cloned into the pETduet vector with a His_6_-MBP-TEV tag at the N terminus of each construct, and expressed by *Escherichia coli* strain BL21(DE3). ^2^H,^13^C,^15^N-labelled Ctc1 OB-A was expressed from M9 minimal media with 0.5 liters of D_2_O, 2 g of ^13^C D-glucose, and 0.5 g of ^15^N ammonium chloride, while ^13^C,^15^N-labelled p50 peptide was expressed from M9 minimal media with ^13^C D-glucose and of ^15^N ammonium chloride. Cultures were grown to mid-log phase at 37°C, induced by the addition of β-D-1-thiogalactopyranoside (IPTG) to a final concentration of 0.5 mM, and incubated at 18°C for an additional 12 h before harvesting by centrifugation. The purification steps for Ctc1 OB-A and p50 peptides were similar to those for *Tetrahymena* PolαPrim described above. Briefly, cells were re-suspended in buffer A, lysed by sonication and centrifuged at 4,500×*g* for 30 min. The supernatant was loaded onto a 5-ml HisTrap HP column pre-equilibrated in buffer A. The column was washed with buffer A and the protein was eluted with buffer B. The protein was digested overnight with 0.2 mg/ml TEV protease and buffer-exchanged to buffer A. The digest was applied onto a 5-ml HisTrap HP column pre-equilibrated in buffer A. The flow-through was concentrated and further purified on an Superdex 75 gel filtration column pre-equilibrated in buffer C. Fractions containing pure protein were pooled, buffer exchanged into protein NMR buffer [20 mM Tris (pH 7.5), 50 mM NaCl, 1 mM TCEP, 3 mM NaN_3_, and 8% D_2_O], and concentrated to 0.5-0.8 mM for NMR studies.

### NMR spectroscopy and data processing

NMR experiments were performed at 298 K on 800 and 600 MHz Bruker spectrometers equipped with HCN cryoprobes. The backbone assignments of Ctc1 OB-A were obtained using the TROSY-type HNCACB, HN(CO)CACB, HNCA, HN(CO)CA, HNCO, and HN(CA)CO spectra collected on an 800 MHz Bruker instrument with ^2^H,^13^C,^15^N-labeled Ctc1 OB-A. Conventional triple resonance backbone assignment experiments (HNCACB, CBCA(CO)NH, HNCA, HN(CO)CA) were used for p50 peptide (228-250) backbone assignments. Spectra were collected and processed using Topspin 4.1, and analyzed with CARA (http://cara.nmr.ch) and NMRFAM Sparky^56^ to interactively obtain sequence-specific resonance assignments. NOE peak lists were automatically generated by Atnos^57^, and assigned by CYANA 3.98 (ref^58^). Secondary structure of Ctc1 OB-A and p50 peptide (228-250) were predicted by software TALOS+ (ref^44^). The model of p50 peptide (228-250) was generated by CS-Rosetta^45^. To investigate the interaction between Ctc1 OB-A and p50 peptides, ^1^H-^15^N HSQC spectra of titration of unlabeled p50 peptides with labeled Ctc1 OB-A and unlabeled Ctc1 OB-A with labeled p50 peptides were obtained, respectively. Chemical shift mapping was analyzed by comparing the apo and bound-form HSQC spectra. The chemical shift perturbation value for each residue was calculated as 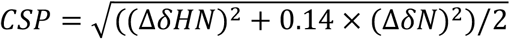 (ref^59^). The backbone assignments of ^2^H,^13^C,^15^N-labeled Ctc1 OB-A in the presence of unlabeled p50 peptide (228-250) were obtained using the same TROSY-type spectra as listed above, while the backbone assignment of ^13^C,^15^N-labeled p50 peptide (228-250) in the presence of unlabeled Ctc1 OB-A were obtained using conventional triple resonance spectra.

### Telomerase sample preparation

*Tetrahymena* telomerase holoenzyme was expressed and purified as described previously^40,60^. To prepare the telomerase–PolαPrim complex sample, 0.5 μM of purified POLA1–POLA2–PRIM1–PRIM2 was incubated with telomerase holoenzyme at the anti-Flag M2 affinity gel (Sigma) step overnight at 4°C, in the presence of excess d(GTTGGG)_10_ primer. Excess DNA and PolαPrim were removed with wash buffer [20 mM HEPES (pH 8.0), 50 mM NaCl, 1 mM MgCl_2_, 1 mM TCEP, 10% (v/v) glycerol, 0.1% (v/v) IGEPAL CA-630] and the final product was eluted using a small volume (30-50 μL) of elution buffer [20 mM HEPES (pH 8.0), 50 mM NaCl, 1 mM MgCl_2_, 1 mM TCEP, and 0.1% (v/v) IGEPAL CA-630] supplemented with 1 mg/mL 3× FLAG peptide.

### Cryo-EM specimen preparation and data collection

For telomerase–PolαPrim complex, 3 μL of the purified sample was applied to glow-discharged lacey carbon grids with a supporting ultrathin carbon film (Ted Pella). The grids were then blotted with filter paper and flash-frozen in liquid ethane using an FEI Vitrobot Mark IV at 10°C and 100% humidity. Cryo-EM grids of PolαPrim were prepared similarly with Quantifoil 200 mesh R2/1 grids. Cryo-EM grids were loaded into a ThermoFisher Titan Krios electron microscope operated at 300 kV for automated data collection using SerialEM^61^. Movies of dose-fractionated frames were acquired with a Gatan K3 direct electron detector in super-resolution mode at a pixel size of 0.55 Å on the sample level. A Gatan Imaging Filter (GIF) was inserted between the electron microscope and the K3 camera and operated at zero-loss mode with the slit width of 20 eV. The microscope was carefully aligned prior to each imaging session and parallel beam was optimized using coma-free alignment in SerialEM. The total dose rate on the sample was set to ~55 electrons/Å^2^, which was fractionated into 50 frames with 0.06 s exposure time for each frame. For telomerase–PolαPrim, 36,716 movies were collected in two separate imaging sessions with the same batch of cryo-EM grids. For PolαPrim, 7,120 movies were collected in a single imaging sessions.

### Cryo-EM data processing

Cryo-EM data processing workflows are outlined in Extended Data Fig. 1, 6 and 7 for the structure determination of *Tt*CST in telomerase, PolαPrim-bound *Tt*CST in telomerase, and PolαPrim alone, respectively. All steps described below were performed with RELION 3.1 (ref^62^) unless otherwise indicated.

To determine *Tt*CST structure, telomerase particles selected from three published datasets (the T3D2, T4D4 and T5D5 datasets as detailed in ref^40^) were combined, resulting in over 2.5 million good particles (Extended Data Fig. 1a). Refinement of these particles without a mask generated a reconstruction with only weak density for *Tt*CST, which confirms the multiple orientation of *Tt*CST relative to the rest of telomerase holoenzyme^23^. To separate particles with *Tt*CST at different positions, an alignment-free 3D classification was performed using a spherical mask covering the *Tt*CST region (mask1). Particles from classes with *Tt*CST at similar positions were grouped together and refined, resulting in three reconstructions (P1, P2 and P3 in Extended Data Fig. 1a), among which P1 has the best density and the largest number of particles. To improve the overall density of P1, we performed another round of 3D classification with local angular search (RELION options: --sigma_ang 8 --healpix_order 4). A soft mask (mask2) was used in this step to exclude the flexible p65. 259,330 particles from the best class were selected for 3D refinement, following by refinement of contrast transfer function (CTF) parameters and Bayesian polishing in RELION. The resulting “shiny” particles were refined to 3.5 Å resolution for the entire telomerase holoenzyme including *Tt*CST (Extended Data Fig. 1b-d). An additional focused 3D classification step was conducted to improve the local resolution of Ctc1 OB-A. The resulting 78,471 particles were refined to 3.8 Å resolution using mask2.

For the newly collected telomerase–PolαPrim and PolαPrim datasets, dose-fractionated frames of each movie were 2x binned (pixel size of 1.1 Å), aligned for the correction of beam-induced drift, and dose weighted using RELION’s implementation of UCSF MotionCor2 (ref^63^). CTF parameters, including defocus and astigmatism, of each dose-weighted micrograph were determined by CTFFIND4 (ref^64^) within RELION.

Two datasets of telomerase–PolαPrim, one for each data collection session, were initially processed separately (Extended Data Fig. 6a). Particles picked from 2,000 representative micrographs using template-free auto-picking in RELION were screened by 2D classification, and the best particles were selected to train a particle detection model in Topaz^65^ for subsequent neural-network based particle picking for all micrographs. After several rounds of 2D and 3D classifications as detailed in Extended Data Fig. 6a, good particles selected from two datasets were combined, resulting in over 1.6 million particles. Refinement of these particles without using any mask generated a reconstruction with only weak density for *Tt*CST– PolαPrim, suggesting that *Tt*CST–PolαPrim also has multiple orientations relative to the rest of telomerase holoenzyme, including telomerase core RNP, TEB heterotrimer and p50, as previously observed for the *Tt*CST dataset (Extended Data Fig. 1a). Therefore, these two parts were processed separately in the following steps. For telomerase core RNP–TEB–p50, a soft mask was used to exclude the dynamic *Tt*CST–PolαPrim during 3D refinement, which resulted in a 3.1 Å resolution reconstruction. After an additional round of focused classification with local angular search in 8° (RELION options: --sigma_ang 8 --healpix_order 4), 539,078 particles from the best class were selected and refined to 2.9 Å resolution (Extended Data Fig. 6b, c). For *Tt*CST–PolαPrim, three rounds of alignment-free 3D classification with an optimized regularization parameter (RELION option: --tau2_fudge 16) were performed in parallel using a spherical mask covering the *Tt*CST–PolαPrim region (mask2). 427,158 particles from classes with interpretable *Tt*CST–PolαPrim densities were combined after removing duplicates. Refinement of these particles generated a reconstruction with clear *Tt*CST–PolαPrim density. Then, we shifted the center of each particle to *Tt*CST–PolαPrim and performed signal subtraction using mask4 to only keep the signal from *Tt*CST–PolαPrim. After two rounds of 3D classification using mask4, 142,912 particles were selected and refined using the same mask, which resulted in a 4.2 Å resolution reconstruction for *Tt*CST–PolαPrim (Extended Data Fig. 6b, d). These particles were back projected to original particles without signal-subtraction and refined to 4.4 Å resolution for the entire complex including both telomerase and *Tt*CST– PolαPrim.

For the PolαPrim dataset (Extended Data Fig. 7), particle picking was conducted using Topaz^65^ in a similar way as described above for the PolαPrim bound telomerase datasets. After two rounds of 2D classification, 1.3 million particles were selected and refined using an initial model generated by cryoSPARC^66^. The resulting cryo-EM map has a “head” that is the catalytic core domain of POLA1 and a bow-tie-shaped “body” that contains POLA2, PRIM1, PRIM2_N_ and the C-terminal domain of POLA1. The “head” and the “body” have multiple orientations relative to each other as indicated by their low-resolution densities in the 3D reconstruction and the 2D classification results (Extended Data Fig. 7b). Focused refinement of the “head” didn’t work well due to its small size, but focused refinement of the “body” generated a map with clear secondary structure features. After one round of 3D classification using the same mask for the “body”, 264,498 particles were selected and refined to 4.5 Å resolution. During the 3D classification step, a notable hinge movement was observed within the “body”. Due to that, we further refined the two halves of the “body” individually and obtained a 4.0 Å resolution reconstruction for POLA2–POLA1_CTD_–PRIM2_N_ and a 4.3 Å resolution reconstruction for PRIM2_N_–PRIM1 (Extended Data Fig. 7c-e).

All cryo-EM maps were sharpened with a negative B-factor and low-pass filtered to the stated resolution using the *relion_postprocess* program in RELION. Local resolution evaluations were determined by ResMap^67^ with two independently refined half-maps. Data collection and processing statistics are given in Extended Data Table 1.

### Model building and refinement

For the modeling of *Tt*CST, crystal structures of *Tetrahymena* Ten1–Stn1-OB^23,24^ (PDB 5DOI and 5DFM) and cryo-EM structure of *Tetrahymena* telomerase RNP–TEB–p50^40^ (PDB 7LMA) were initially rigid-body fitted into the 3.5 Å resolution cryo-EM map (Extended Data Fig. 1c) using UCSF Chimera^68^, and refined manually in COOT^69^. Density of Ctc1 was traced from its signature C-terminal α helix, and models of OB-C and OB-B were built *de novo* against the density in COOT. Visible densities of amino acid residues with bulky side chains, such as Phe, Tyr and Trp were used as guidance for sequence assignment (Extended Data Fig. 1f). For Ctc1 OB-A, an initial model was built against the cryo-EM density and refined manually with structure information obtained from NMR (Extended Data Fig. 3). Briefly, secondary structure information obtained from TALOS+ (ref^44^) was used to define the boundaries of β strands within the OB (Extended Data Fig. 3d), and 105 inter-β-strand NOE restraints were used to refine the relative position of the β strands (Extended Data Fig. 3e). Lastly, p50 residues 184-208 following the C-terminus of the previous p50 OB model (PDB 7LMA) were built into the cryo-EM map adjacent to Ctc1 OB-A (Extended Data Fig. 1g).

For the modeling of *Tt*CST–PolαPrim, the *Tt*CST model obtained as described above, crystal structures of Stn1 WH-WH^23,24^ (PDB 5DFN and 5DOK), and a computed model of POLA1 core generated using AlphaFold2 (ref^47^) were initially rigid-body fitted into the 4.2 Å resolution map (Extended Data Fig. 6d) using UCSF Chimera^68^, and refined manually in COOT^69^ (Extended Data Fig. 6f). A segment of sstDNA was built manually against the density in the C-shape cleft of Ctc1 OB-C. The previously reported cryo-EM structure of *Tetrahymena* telomerase RNP–TEB–p50^40^ (PDB 7LMA) was refined against the 2.9 Å resolution map (Extended Data Fig. 6c).

For the modeling of PolαPrim platform (POLA2–POLA1_CTD_–PRIM2_N_–PRIM1), a composite cryo-EM density map was generated using the “combine focused maps” function in Phenix^70^ with two focused refined maps (Extended Data Fig. 7a, e). Computed models of individual subunits generated using AlphaFold2 (ref^47^) were rigid-body fitted into the composite map and refined manually in COOT^69^ (Extended Data Fig. 7g).

All models were refined using Phenix^70^ in real space with secondary structure, Ramachandran, and rotamer restraints. Refinement statistics of the models are summarized in Extended Data Table 1. Model vs map FSC validations were shown in Extended Data Fig. 1e, 6e and 7f. All figures presenting the model were prepared using UCSF ChimeraX^71^.

### Electrophoretic mobility shift assay (EMSA)

*Tetrahymena* CST–p50 and its mutants were expressed and purified from insect cells as described above. Radioactive ^32^P 5’-labeled (GTTGGG)_10_ DNA primer was used for EMSAs. *Tt*CST–p50 proteins were incubated with DNA primer in 10 μl EMSA buffer [20 mM HEPES-NaOH (pH 8.0), 150 mM NaCl, 2 mM MgCl_2_, 5% glycerol, and 1 mM TCEP] for 1 hour on ice before being loaded onto a 0.5× TBE 0.7% agarose gel. The gel was vacuum dried and exposed to a phosphor imaging screen for overnight. The final image was obtained by scanning the screen on a ImageFX system (Bio-Rad) and then quantified with QuantityOne (Bio-rad).

## Data availability

Cryo-EM density maps have been deposited in the Electron Microscopy Data Bank under accession numbers xxxx (telomerase with CST), xxxx (telomerase with CST–PolαPrim), xxxx (telomerase), xxxx (CST–PolαPrim), xxxx (PolαPrim platform), xxxx (POLA2–POLA1_CTD_–PRIM2_N_) and xxxx (PRIM2_N_–PRIM1). The atomic models have been deposited in the Protein Data Bank under accession codes xxxx (telomerase with CST), xxxx (telomerase), xxxx (CST–PolαPrim) and xxxx (PolαPrim platform). Backbone chemical shifts have been deposited in BMRB under accession codes xxxx (Ctc1 OB-A) and xxxx (p50 peptide 228-250).

## Acknowledgements

This work was supported by grants from NIH R35GM131901 and NSF MCB2016540 to J.F. and NIH R01GM071940 to Z.H.Z. We acknowledge use of instruments at the Electron Imaging Center for Nanomachines supported by UCLA and instrumentation grants from NIH (1S10OD018111 and U24GM116792) and NSF (DBI-1338135 and DMR-1548924). The UCLA-DOE NMR core facility is supported in part by US Department of Energy grant DE-AC02-06CH11357 and NIH instrumentation grants S10OD016336 and S10OD025073.

## Author Contributions

Y.H. and H.S prepared and checked EM samples; Y.H collected and analyzed cryo-EM data; Y.H. and H.S. built the models; H.C. expressed and purified samples for NMR analysis; H.C., Y.W., and L.S. collected and analyzed NMR data; B.L. and Y.H. conducted assays; Z.H.Z. supervised cryo-EM data collection and processing; J.F. supervised all aspects of the project; Y.H. and J.F. made figures and wrote the manuscript, with input from H.S. and Y.W.

---

**Extended Data Fig. 1:**
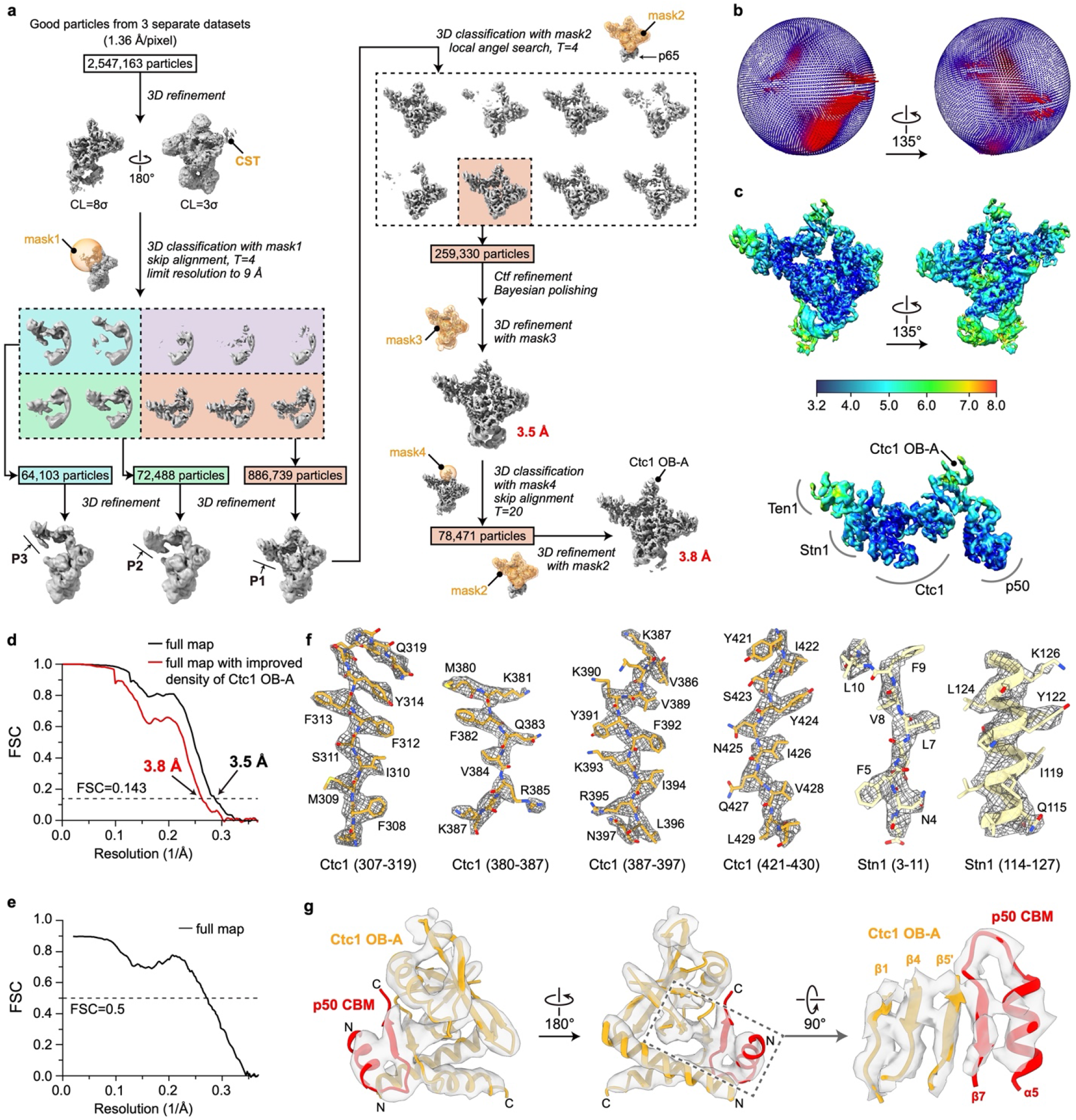
Cryo-EM data processing workflow of *Tt*CST in telomerase holoenzyme and evaluations of the final reconstructions. **a,** Data processing workflow (detailed in Methods). **b,** Euler angle distributions of particles used for the final 3.5 Å resolution reconstruction. **c,** Local resolution evaluation of the 3.5 Å resolution reconstruction shown for overall map (upper) and for the *Tt*CST–p50 region (lower). **d,** Plot of the Fourier shell correlation (FSC) as a function of the spatial frequency demonstrating the resolutions of final reconstructions. **e,** FSC coefficients as a function of spatial frequency between model and cryo-EM density map. **f,** Representative cryo-EM densities (gray and mesh) encasing the related atomic models (colored sticks and ribbons). **g,** Superimposition of cryo-EM densities (low-pass filtered to 5 Å) and model of Ctc1 OB-A in complex with p50 CBM.

**Extended Data Fig. 2:**
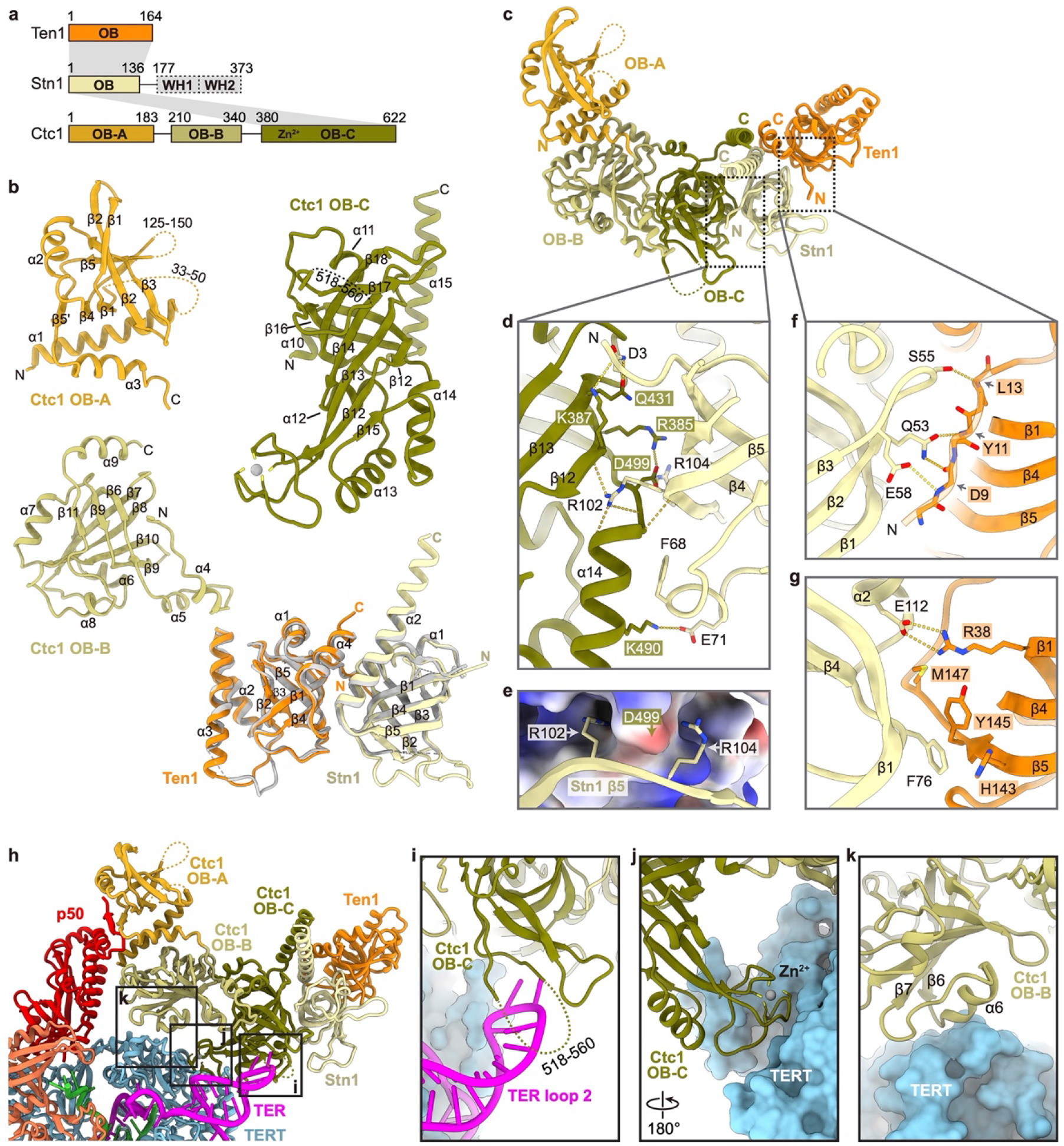
Structural details of *Tt*CST. **a,** Domain organization of *Tt*CST subunits. Invisible regions in the cryo-EM map are shown as dashed boxes. Intermolecular interactions are indicated as gray shading. **b,** Ribbon diagrams of *Tt*CST subunits/domains with secondary structure elements labelled. Unmodeled regions are shown as dashed lines. Crystal structure of Ten1–Stn1 OB (PDB 5DOI) is shown in gray and overlayed with the cryo-EM structure for comparison. **c,** Ribbon representation of *Tt*CST structure with individual OB domain colored as indicated. **d,** Zoom-in view of the interface between Ctc1 OB-C and Stn1 OB. Salt bridge and hydrogen-bonding interactions are shown as dashed yellow lines. **e,** Two arginine sidechains on Stn1 OB (ribbon) clamp D499 on Ctc1 OB-C (electrostatic surface). **f, g,** Detailed interactions between Stn1 OB and Ten1. **h,** Overall view of the interface between *Tt*CST and TERT–TER. **i-k,** Zoom-in views of interactions between *Tt*CST and TERT–TER as indicated in dashed boxes in **h**. These interactions stabilize *Tt*CST in the predominant conformation.

**Extended Data Fig. 3:**
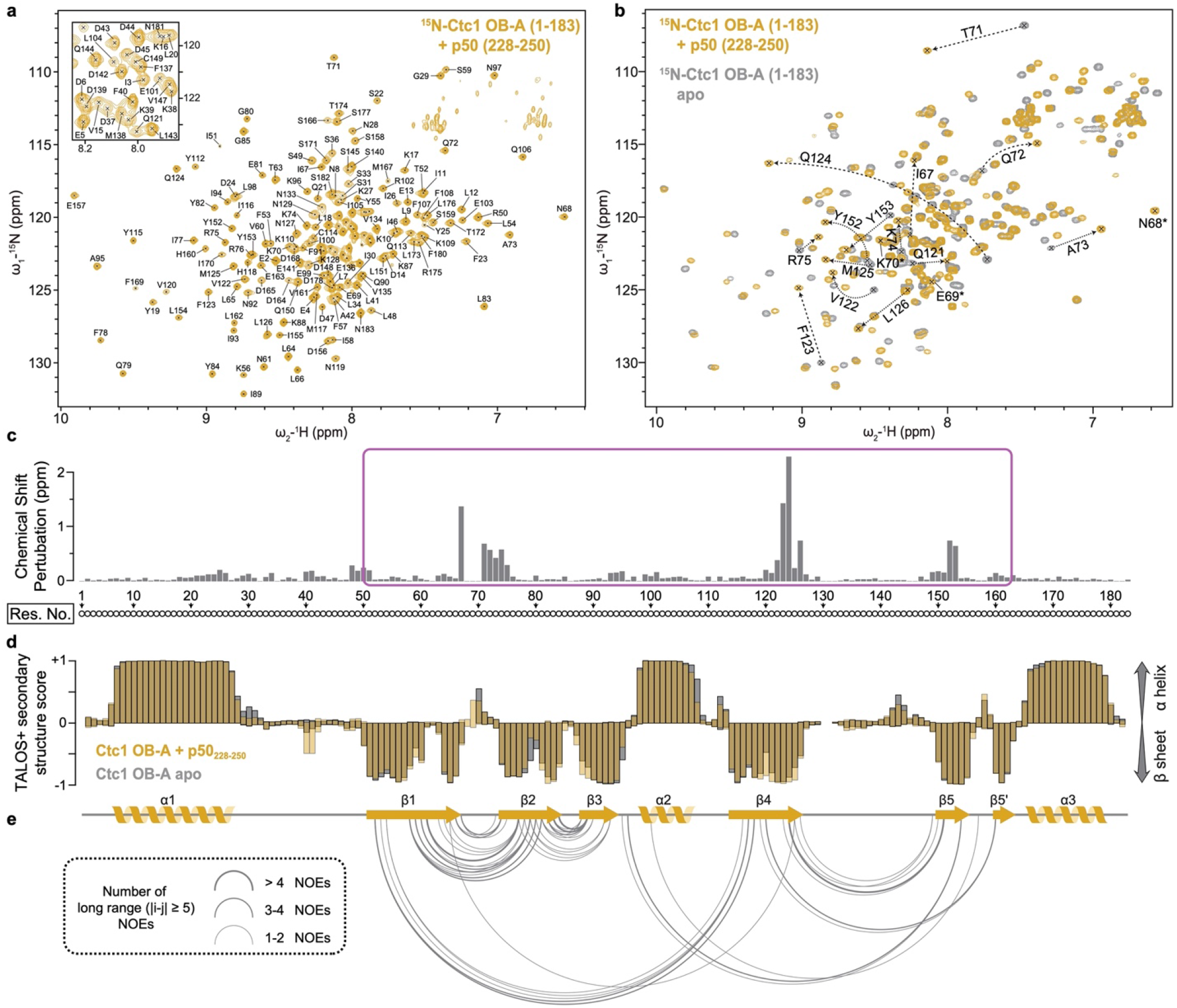
NMR spectra and structural study of Ctc1 OB-A with p50 peptide. **a,** Assigned ^1^H-^15^N HSQC spectrum of ^15^N-labelled Ctc1 OB-A (residues 1-183) in the presence of unlabeled p50 peptide (residues 228-250). Inset shows the expanded central region of the spectrum. **b,** Superimposed ^1^H-^15^N HSQC spectra of ^15^N-labelled Ctc1 OB-A in the presence (yellow) and absence (gray) of unlabeled p50 peptide. Signals from the same residues with chemical shift differences of >0.25 ppm are connected by dashed arrows. Signals from residues 68-70 that only appear in the presence of p50 peptide are labeled with asterisks. **c,** Chemical shift perturbation (CSP) index of ^15^N-labelled Ctc1 OB-A upon binding p50 peptide. Magenta box indicates the region that is shown in Fig. 2d. **d,** Chemical-shift-based secondary structure score of Ctc1 OB-A in the presence (yellow) and absence (gray) of p50 peptide. The scores are determined using TALOS+ (ref^44^). Top and bottom edges of each bar represent the probabilities of each residue assigned to be α helix and β sheet, respectively. The secondary structure of Ctc1 OB-A observed in the cryo-EM structure is shown below for comparison. **e,** Plot of long range (greater than 5 residues) ^1^H-^1^H NOE restraints observed within Ctc1 OB-A. Residues with pairwise NOE restraint(s) are connected by a link. Thickness of each link is coded as indicated based on the number of NOE restraints between the two connected residues.

**Extended Data Fig. 4:**
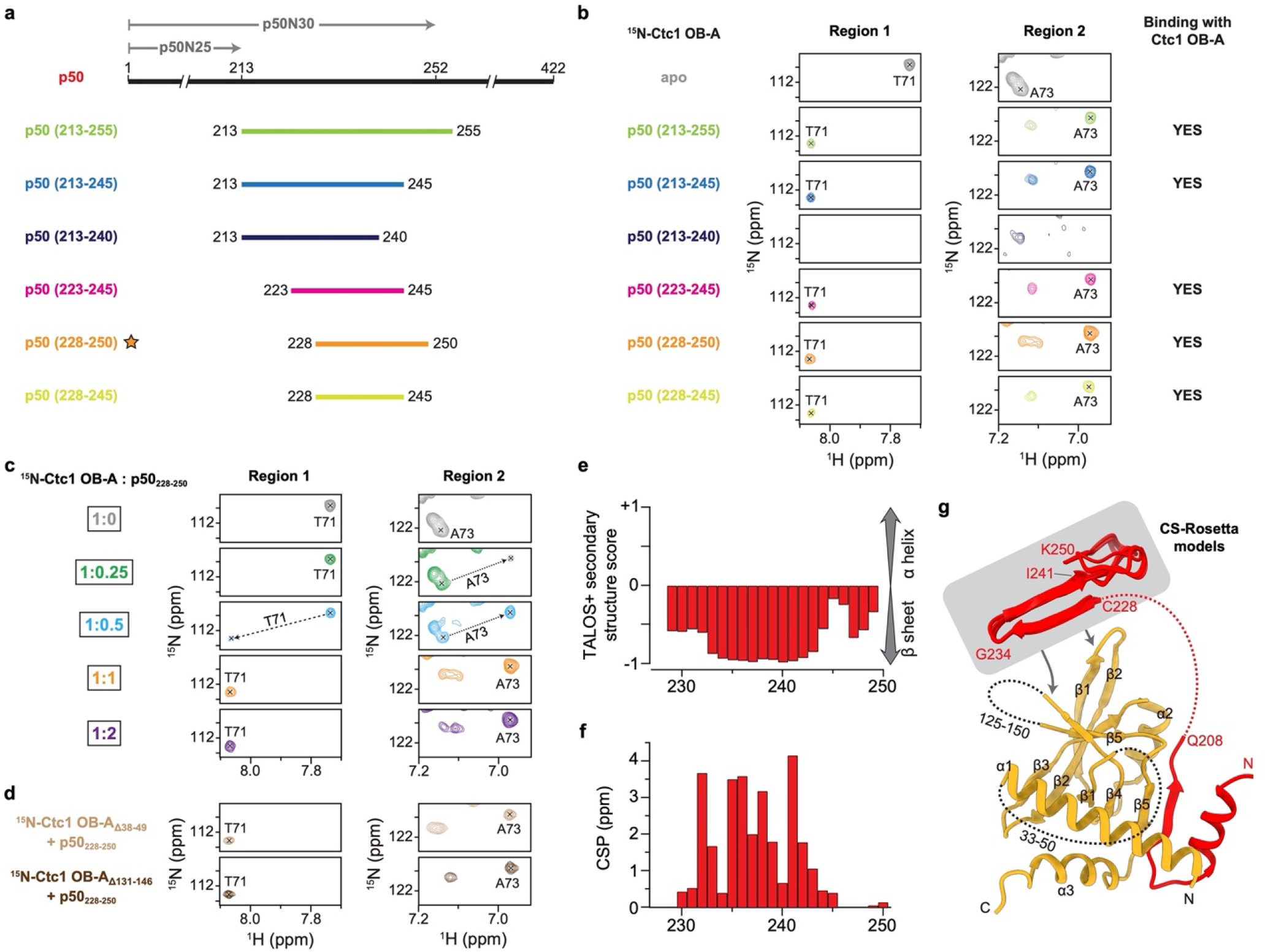
Identifying the “invisible” interface between Ctc1 OB-A and p50 peptide using NMR methods. **a,** Schematic diagram of p50 and constructs of p50 peptide. The N-terminal 30 kDa and 25 kDa fragments of p50 are labeled as p50N30 and p50N25, respectively. Previous biochemical study showed that p50N30 could bind Ctc1, whereas p50N25 could not^43^. The cryo-EM structure of p50 ends at residue 208 (Fig. 2b). Based on these facts, a series of p50 peptides in the range of residues 213-255 were designed to explore additional interface between p50 and Ctc1 OB-A that are “invisible” in the cryo-EM structure. **b,** NMR binding study of p50 peptides with Ctc1 OB-A. Two regions of ^1^H-^15^N HSQC spectra of ^15^N-labelled Ctc1 OB-A in the absence (apo) and presence of unlabeled p50 peptides were shown. Chemical shifts of T71 and A73 were chosen to illustrate the binding process in this and the following panels **c** and **d**. p50_228-250_ peptide is determined to be the optimal construct and was used for other NMR studies presented in this paper. **c,** Titration series of p50 peptide into ^15^N-labelled Ctc1 OB-A. The binding is in the slow exchange regime and saturated at 1:1 stoichiometry. **d,** Truncations of two unstructured loops (residues 38-49 and 131-146) of Ctc1 OB-A individually have no effect on its binding with p50 peptide. **e,** Secondary structure score of p50_228-250_ in the presence of Ctc1 OB-A. **f,** CSP index of p50 peptide upon binding Ctc1 OB-A. ^1^H-^15^N HSQC spectra shown in Fig. 2c were used for the CSP calculation. **g,** Model of the interactions between Ctc1 OB-A and p50. CS-Rosetta models of p50_228-250_ are shown in gray box with arrows pointing to the binding surface on Ctc1 OB-A. Unstructured linkers are shown as dashed lines.

**Extended Data Fig. 5:**
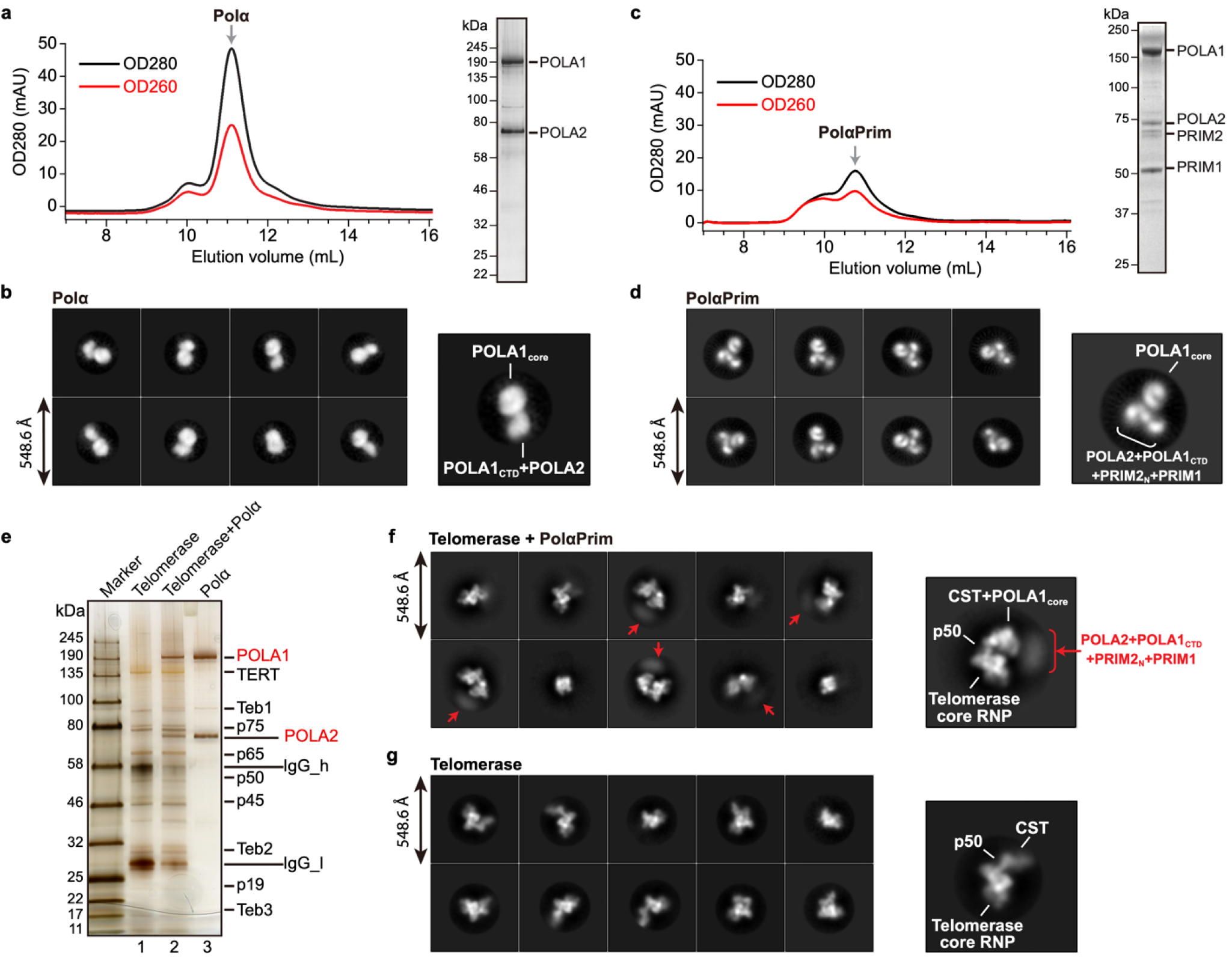
Characterization of purified *Tetrahymena* PolαPrim samples and their assembly with telomerase holoenzyme. **a,** Size-exclusion chromatography (SEC) profile (left) and SDS-PAGE gel (right) of Polα. **b,** Representative 2D-class averages of Polα particles obtained from negative-stain EM images. **c,** SEC profile (left) and SDS-PAGE gel (right) of PolαPrim. **d,** Representative 2D-class averages of PolαPrim particles obtained from negative-stain EM images. **e,** SDS-PAGE gel of affinity purified telomerase–Polα complex (lane 2) shows that Polα can bind telomerase in the absence of Primase. Telomerase (lane 1) and Polα (lane 3) samples were loaded on the same gel for comparison. **f,** Representative 2D-class averages of affinity purified telomerase– PolαPrim obtained from negative-stain EM images. Densities are assigned based on the cryo-EM structure (Fig. 3a) obtained with the same batch of sample. Smeared densities (red arrows) are observed near POLA1_core_ in several classes, so we were able to assign them to the PolαPrim platform, which comprises POLA2, POLA1_CTD_, PRIM2_N_, and PRIM1. **g,** Representative 2D-class averages of telomerase particles shown for comparison with **f**.

**Extended Data Fig. 6:**
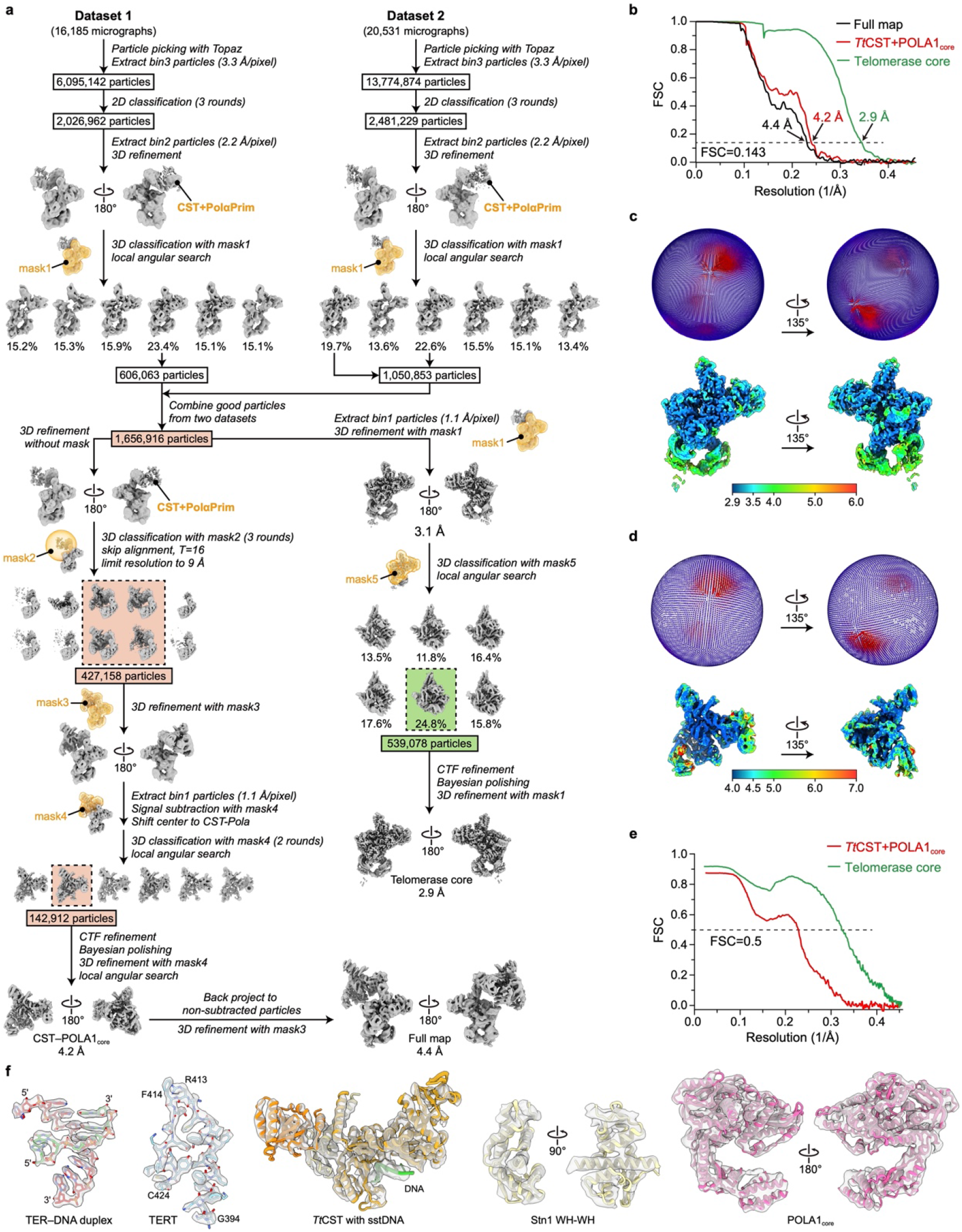
Cryo-EM structure determination of *Tetrahymena* telomerase–PolαPrim complex. **a,** Data processing workflow (detailed in methods). **b,** Resolution of final reconstructions determined by gold-standard FSC at the 0.143 criterion. **c,** Particle distribution (upper) and local resolution evaluation (lower) of the 2.9 Å resolution reconstruction of telomerase core. **d,** Particle distribution (upper) and local resolution evaluation (lower) of the 4.2 Å resolution reconstruction of *Tt*CST–POLA1_core_. **e,** FSC curves for refined models versus the corresponding cryo-EM density maps. **f,** Representative cryo-EM densities (transparency surface) encasing the related atomic models (color sticks and ribbons).

**Extended Data Fig. 7:**
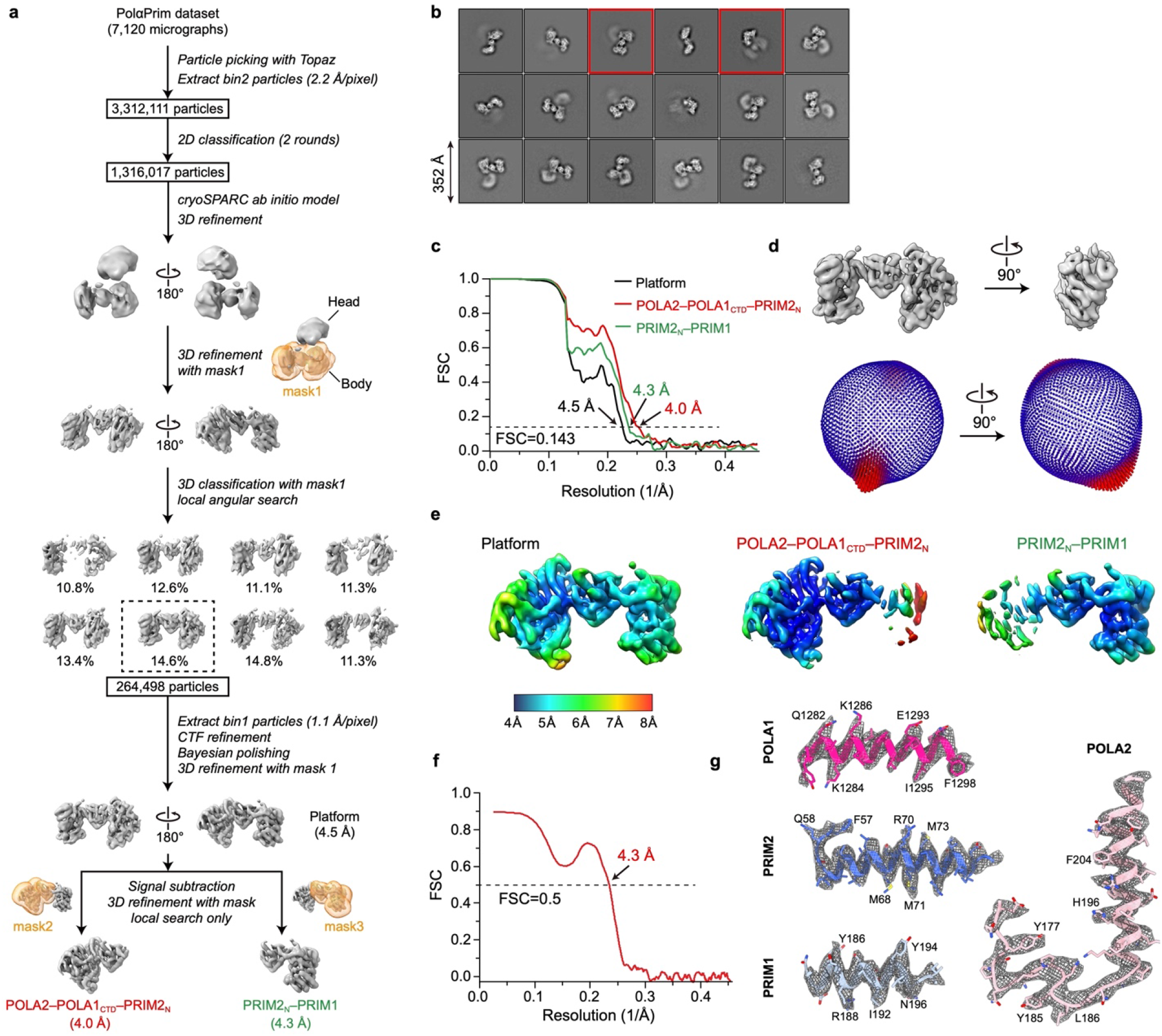
Cryo-EM structure determination of PolαPrim. **a,** Data processing workflow (detailed in methods). **b,** Representative 2D-class averages of PolαPrim particles obtained from cryo-EM images. The two classes shown in Fig. 3e are labeled with red boxes. **c,** Resolution of final reconstructions determined by gold-standard FSC at the 0.143 criterion. **d,** Euler angle distributions of particles used for the final reconstructions. **e,** Local resolution evaluations of the final reconstructions. **f,** FSC curve for the refined model of PolαPrim platform versus the composite density map used for modeling. The composite map is generated using Phenix^70^ with two focused refined maps (detailed in methods). **g,** Representative cryo-EM densities (gray mesh) encasing the related atomic models (colored sticks and ribbons).

**Extended Data Fig. 8:**
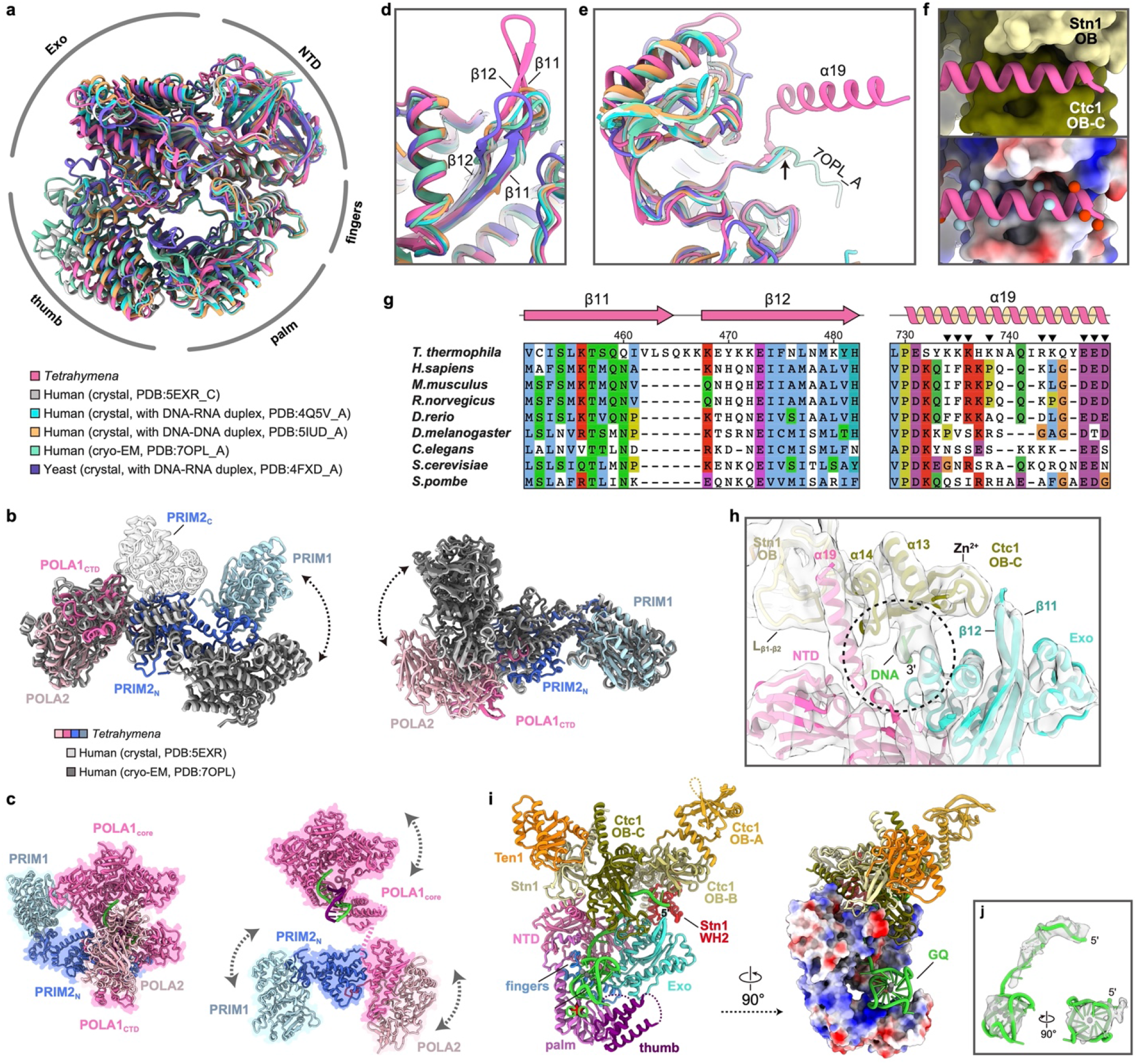
Structural conservation of *Tetrahymena* PolαPrim. **a,** Superposition of *Tetrahymena* (*Tt*), human and yeast POLA1_core_ structures shown in an overall view. **b,** Structural comparison of PolαPrim platform of *Tt* and human. The structures were superposed based on POLA2– POLA1_CTD_ (left) or PRIM1 (right). Arrows indicate dynamics of the unaligned regions. PRIM2_C_ in human structures are shown as transparent ribbons. PRIM2_C_ is not observed in the *Tt* structure. **c,** Structures of PolαPrim in an autoinhibited conformation (left, modeled based on PDB 5EXR) and an active conformation (right, modeled based on a low-resolution cryo-EM map in Extended Data Fig. 7a). The DNA-DNA duplexes on POLA1_core_ were modeled based on PDB 5IUD. In the autoinhibited conformation (left), the active site on POLA1_core_ is sterically blocked by POLA1_CTD_ and POLA2 for DNA entry. In the active conformation (right), dynamics of subunits are indicated with arrows. **d-e,** Superposition of *Tt*, human and yeast POLA1_core_ structures for the regions that are on the interface with *Tt*CST. Conserved domains/motifs are labeled as indicated. The β11-β12 hairpin in *Tt* POLA1_core_ is longer than those in human and yeast (d). The α19 is structured only in *Tt* POLA1 when binding *Tt*CST (e). **f,** Zoom-in views of the interface between *Tt* POLA1_core_ α19 (ribbon) and *Tt*CST (surface/electrostatic surface). In the lower panel, locations of positively and negatively charged residues on α19 are indicated using blue and red balls, respectively. **g,** Sequence conservation analysis of the β11-β12 hairpin and α19 of POLA1. Charged residues on α19 are indicated with black arrows. **h,** Zoom-in view of the interface between POLA1_core_ and *Tt*CST with sstDNA. Cryo-EM density are shown as transparent surface. The template entry port formed by POLA1_core_ NTD and Exo and Ctc1 OB-C is indicated by a cycle. **i,** Path of sstDNA in the cryo-EM structure of *Tt* telomerase–PolαPrim. The sstDNA binds in the C-shape cleft of Ctc1 OB-C with its 5’ side, while its 3’ side passes through the template entry port to the active site of POLA1_core_ (right). A G-quadruplex (GQ) formed by four *Tt* telomere repeats (modeled based on PDB 7JKU) is observed on a positively charged DNA binding surface of POLA1_core_ between the palm and thumb (middle). **j,** Superimposition of the GQ structure and cryo-EM density. Weak density of sstDNA can be observed connecting the sstDNA on Ctc1 OB-C to the GQ.

**Extended Data Fig. 9:**
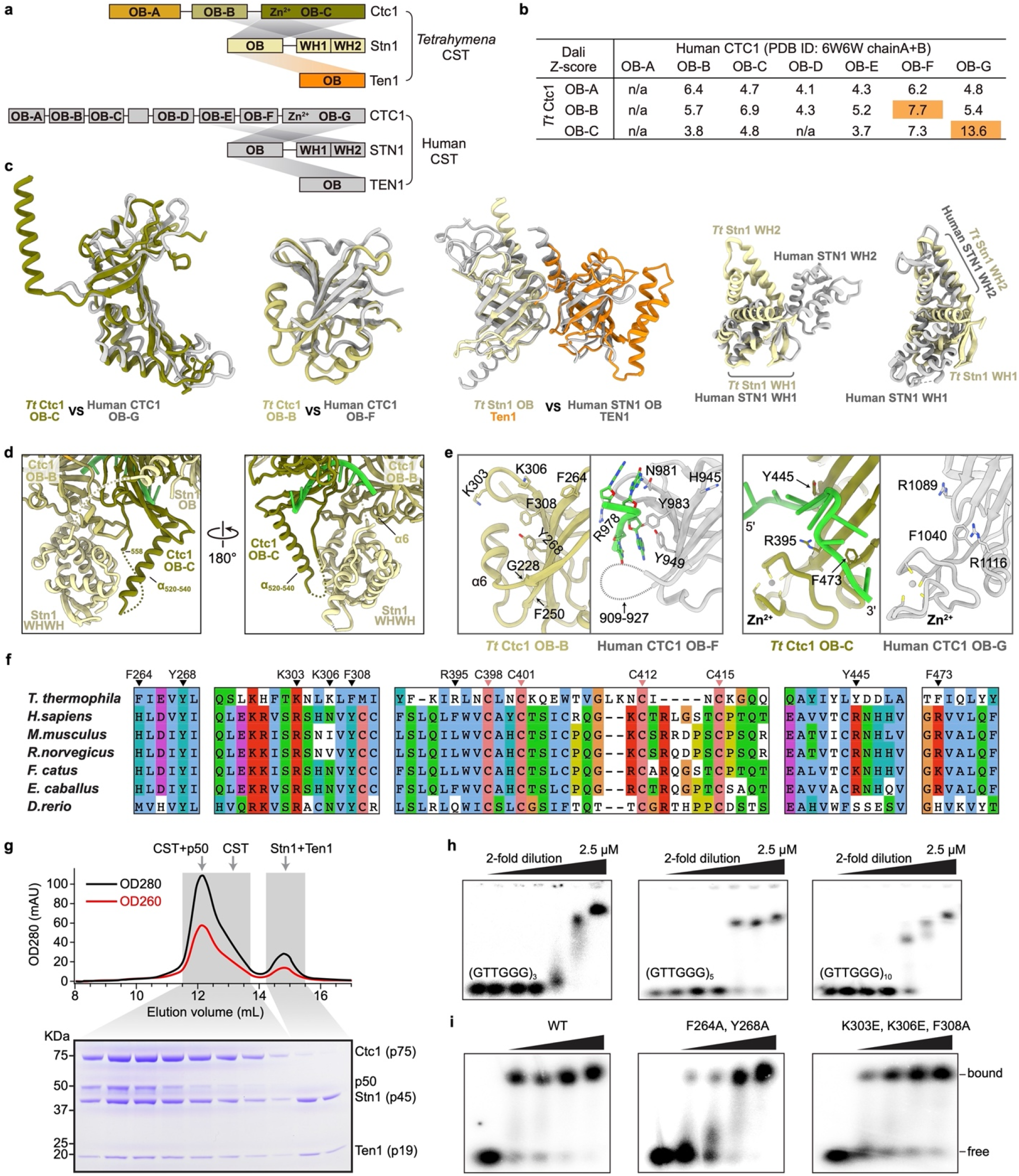
Comparison of *Tt*CST and hCST. **a,** Domain diagrams of *Tt*CST and hCST. **b,** Structural homology analysis of individual OB domains of *Tt* Ctc1 (OB-A to -C) and human CTC1 (OB-A to -G) using the Dali sever^54^. Based on the resulted pairwise Z-scores, *Tt* Ctc1 OB-B and OB-C are identified as homologs of human CTC1 OB-F and OB-G, respectively. **c,** Structural comparison of individual domains from *Tt*CST (color) with corresponding domains from hCST (gray). Structures of WH-WH domains of *Tt* Stn1 and human STN1 were superposed based on WH1 or WH2 domain. The relative orientation of the two WH domains is different between *Tt* and human. **d,** The interface between Stn1 WH-WH and Ctc1 in the cryo-EM structure of *Tt* telomerase–PolαPrim. An previously unstructured loop of Ctc1 OB-C (Extended Data Fig. 2b) partially forms an α helix (α_520-540_) and contributes to the interface with Stn1 WH-WH. **e,** Comparison of DNA binding sites on *Tt* Ctc1 (color) and human CTC1 (gray). Conserved residues located on the DNA binding interface are shown as sticks. In the decameric structure of hCST (PDB 6W6W), sstDNA primarily binds on CTC1 OB-F. However, in *Tt*CST, the equivalent sstDNA binding site on OB-B is partially occluded by a helix (α6) that is part of an unstructured loop in hCTC1 OB-F. The helix α6 abuts TERT in TtCST without PolαPrim (Extended Data Fig. 2k) and Stn1 WH2 when PolαPrim is bound (panel d). **f,** Sequence conservation analysis of Ctc1 residues on the DNA binding interface. Residues shown in panel e are indicated with black arrows. Conserved cysteines in the Zn-ribbon motifs are indicated with pink arrows. **g,** SEC profile and SDS-PAGE gel of *Tt*CST–p50 co-expressed in Sf9 cells. Gel samples are from the peak fractions of the SEC profile as indicated. **h,** EMSA of purified wild-type *Tt*CST with sstDNA of 3, 5 or 10 repeats. **i,** Substitution of *Tt* Ctc1 OB-B conserved residues Y264A/Y268A or K303E/K306E/F308A substantially decreases d(GTTGGG)_5_ binding, as indicated by EMSAs. These results suggest that the binding site on *Tt* Ctc1 OB-B may be accessible to sstDNA in free *Tt*CST where neither TERT nor Stn1 WH-WH stabilize helix α6. Wedges indicate two-fold dilution of *Tt*CST starting at 2.5 μM. The first lane of each gel is a *Tt*CST-free control. Each assay was successfully repeated three times.

**Extended Data Table. 1:**
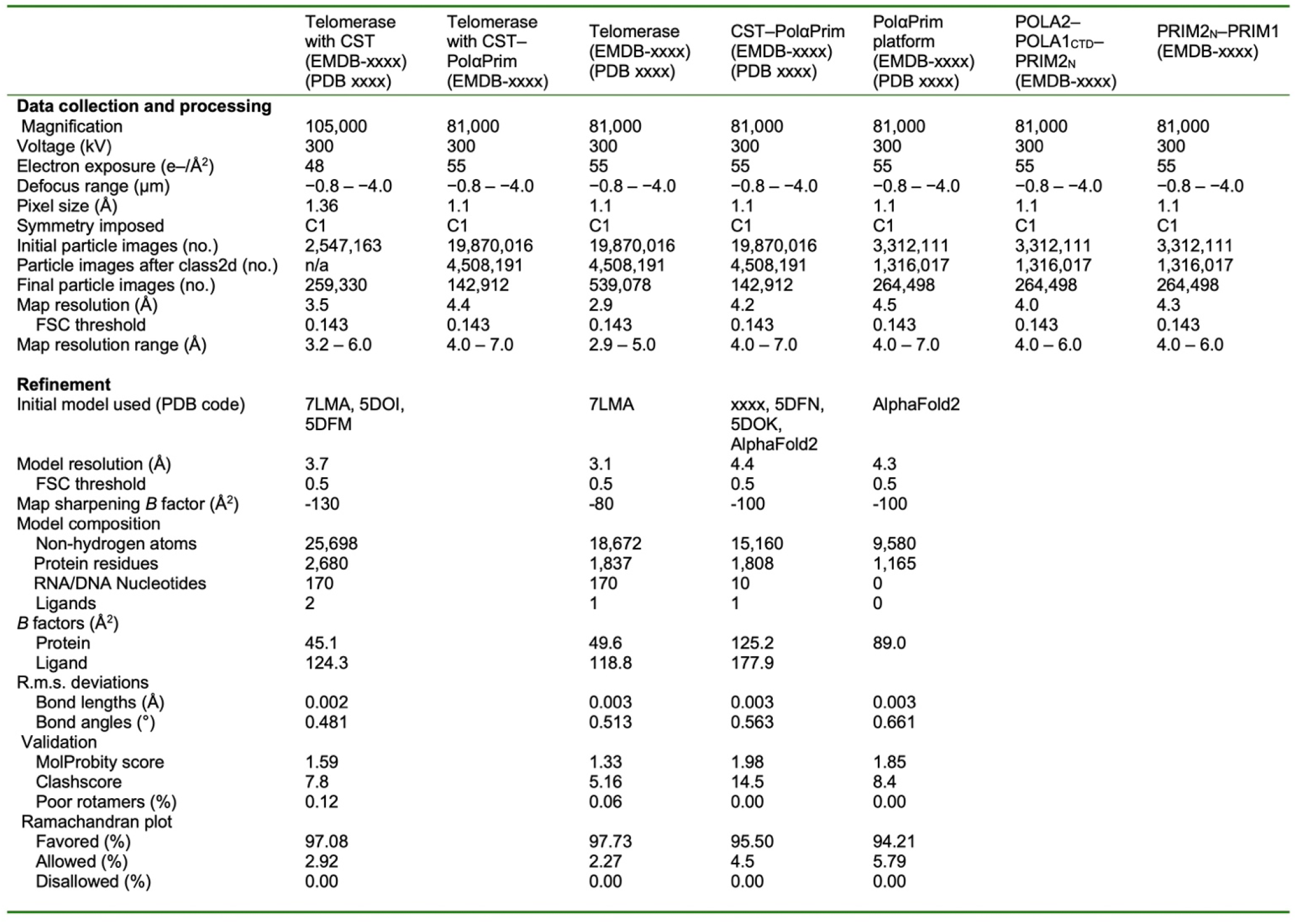
Cryo-EM data collection, refinement and validation statistics.

|  | Telomerase<br>with CST<br>(EMDB-xxxx)<br>(PDB xxxx) | Telomerase<br>with CST-PolaPrim<br>(EMDB-xxxx) | Telomerase<br>(EMDB-xxxx)<br>(PDB xxxx) | CST-PolaPrim<br>(EMDB-xxxx)<br>(PDB xxxx) | PolaPrim<br>platform<br>(EMDB-xxxx)<br>(PDB xxxx) | POLA2-POLA1<sub>CTD</sub>-<br>PRIM2<sub>N</sub><br>(EMDB-xxxx) | PRIM2<sub>N</sub>-PRIM1<br>(EMDB-xxxx) |
| --- | --- | --- | --- | --- | --- | --- | --- |
| **Data collection and processing** |  |  |  |  |  |  |  |
| Magnification | 105,000 | 81,000 | 81,000 | 81,000 | 81,000 | 81,000 | 81,000 |
| Voltage (kV) | 300 | 300 | 300 | 300 | 300 | 300 | 300 |
| Electron exposure (e-/Å<sup>2</sup>) | 48 | 55 | 55 | 55 | 55 | 55 | 55 |
| Defocus range (µm) | -0.8 – -4.0 | -0.8 – -4.0 | -0.8 – -4.0 | -0.8 – -4.0 | -0.8 – -4.0 | -0.8 – -4.0 | -0.8 – -4.0 |
| Pixel size (Å) | 1.36 | 1.1 | 1.1 | 1.1 | 1.1 | 1.1 | 1.1 |
| Symmetry imposed | C1 | C1 | C1 | C1 | C1 | C1 | C1 |
| Initial particle images (no.) | 2,547,163 | 19,870,016 | 19,870,016 | 19,870,016 | 3,312,111 | 3,312,111 | 3,312,111 |
| Particle images after class2d (no.) | n/a | 4,508,191 | 4,508,191 | 4,508,191 | 1,316,017 | 1,316,017 | 1,316,017 |
| Final particle images (no.) | 259,330 | 142,912 | 539,078 | 142,912 | 264,498 | 264,498 | 264,498 |
| Map resolution (Å) | 3.5 | 4.4 | 2.9 | 4.2 | 4.5 | 4.0 | 4.3 |
| FSC threshold | 0.143 | 0.143 | 0.143 | 0.143 | 0.143 | 0.143 | 0.143 |
| Map resolution range (Å) | 3.2 – 6.0 | 4.0 – 7.0 | 2.9 – 5.0 | 4.0 – 7.0 | 4.0 – 7.0 | 4.0 – 6.0 | 4.0 – 6.0 |
| **Refinement** |  |  |  |  |  |  |  |
| Initial model used (PDB code) | 7LMA, 5DOI,<br>5DFM |  | 7LMA | xxxx, 5DFN,<br>5DOK,<br>AlphaFold2 | AlphaFold2 |  |  |
| Model resolution (Å) | 3.7 |  | 3.1 | 4.4 | 4.3 |  |  |
| FSC threshold | 0.5 |  | 0.5 | 0.5 | 0.5 |  |  |
| Map sharpening B factor (Å<sup>2</sup>) | -130 |  | -80 | -100 | -100 |  |  |
| Model composition |  |  |  |  |  |  |  |
| Non-hydrogen atoms | 25,698 |  | 18,672 | 15,160 | 9,580 |  |  |
| Protein residues | 2,680 |  | 1,837 | 1,808 | 1,165 |  |  |
| RNA/DNA Nucleotides | 170 |  | 170 | 10 | 0 |  |  |
| Ligands | 2 |  | 1 | 1 | 0 |  |  |
| B factors (Å<sup>2</sup>) |  |  |  |  |  |  |  |
| Protein | 45.1 |  | 49.6 | 125.2 | 89.0 |  |  |
| Ligand | 124.3 |  | 118.8 | 177.9 |  |  |  |
| R.m.s. deviations |  |  |  |  |  |  |  |
| Bond lengths (Å) | 0.002 |  | 0.003 | 0.003 | 0.003 |  |  |
| Bond angles (°) | 0.481 |  | 0.513 | 0.563 | 0.661 |  |  |
| Validation |  |  |  |  |  |  |  |
| MolProbity score | 1.59 |  | 1.33 | 1.98 | 1.85 |  |  |
| Clashscore | 7.8 |  | 5.16 | 14.5 | 8.4 |  |  |
| Poor rotamers (%) | 0.12 |  | 0.06 | 0.00 | 0.00 |  |  |
| Ramachandran plot |  |  |  |  |  |  |  |
| Favored (%) | 97.08 |  | 97.73 | 95.50 | 94.21 |  |  |
| Allowed (%) | 2.92 |  | 2.27 | 4.5 | 5.79 |  |  |
| Disallowed (%) | 0.00 |  | 0.00 | 0.00 | 0.00 |  |  |

